# Succinate enhances mitochondrial metabolism and phagocytosis in human airspace monocytes

**DOI:** 10.1101/2025.06.12.659271

**Authors:** Karen Slattery, Aisling Newing, John McGrath, Albert Sanfeliu, Oisín O’Gallchobhair, Lynne Faherty, Emma McNally, Finbar O’Connell, Parthiban Nadarajan, Patrick Mitchel, Seamas C. Donnelly, Suzanne M. Cloonan, Joseph Keane, Laura Gleeson

## Abstract

Airspace macrophages (AM) are crucial to host defence and to maintenance of lung homeostasis, with smoking drastically compromising these functions. Airspace monocytes, precursors of the differentiated AM, are found in increased numbers in the lungs of smokers yet little is known about their metabolic regulation and function. Here, we develop a click chemistry-based single cell analysis platform to characterise human airspace monocytes and AM *ex vivo*, identifying distinct metabolic profiles and a key role for oxidative phosphorylation in supporting phagocytic function. While blood and newly recruited CD93+ airspace monocytes show low mitochondrial dependency, AM rely heavily on oxidative phosphorylation. Acute succinate supplementation enhanced mitochondrial metabolism and phagocytosis in monocytes and promoted their differentiation into highly oxidative macrophages with enhanced function. Succinate emerges as a promising candidate to restore lung immune function, particularly in the smoker’s lung where airspace monocytes are enriched. Overall, we identify mitochondrial metabolism as a key modulator of lung immune function and a target for therapeutic intervention, with potential applications in systemic monocyte-targeted therapies and metabolic preconditioning for adoptive cell therapies.

**One Sentence Summary**: Mitochondrial metabolism regulates phagocytic function in human lung monocytes and macrophages and can be targeted using succinate.

**Graphical Abstract:** 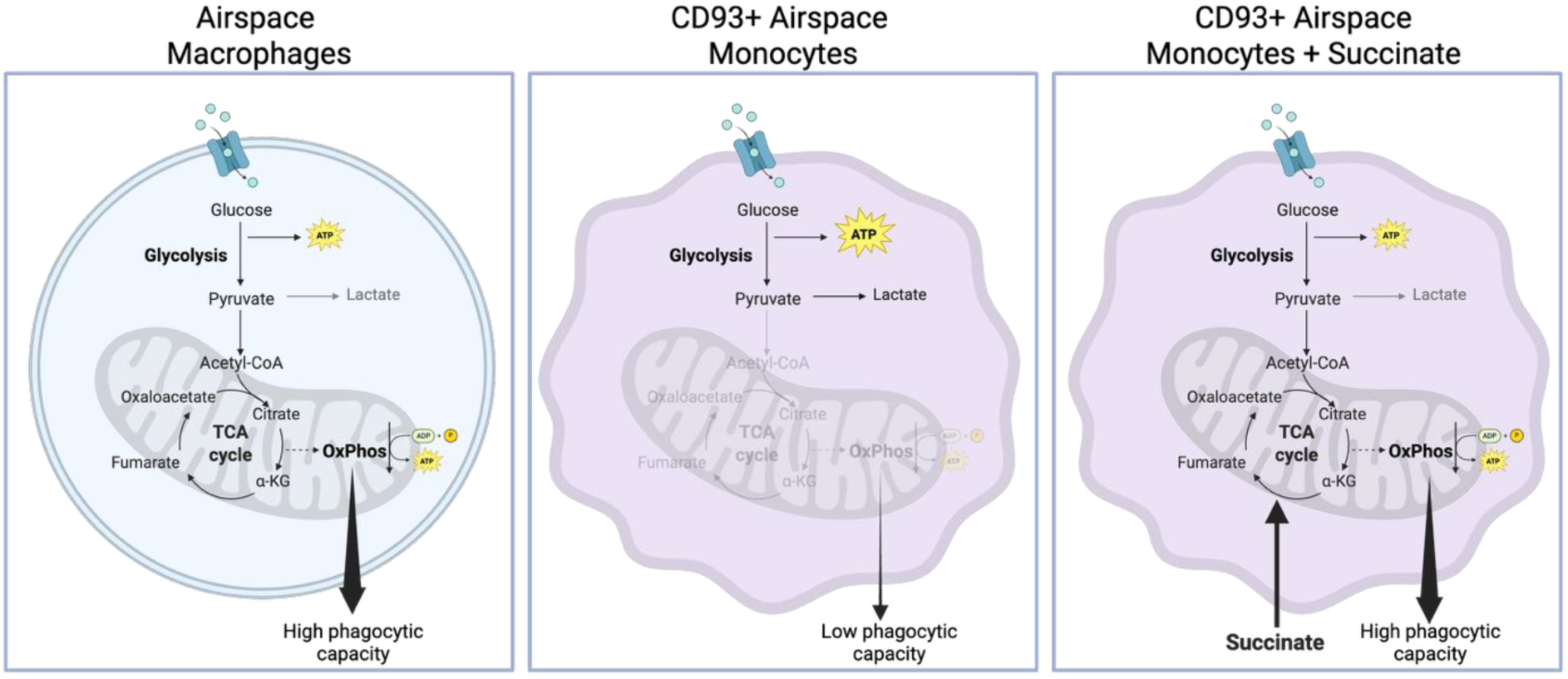

## Introduction

The lungs are a key front line of defence against environmental pollutants and pathogens. Central to this function is the airspace macrophage (AM), the dominant innate immune cell within the lung^1,2^. The AM resides in the alveolar space where it plays a pivotal role in the phagocytosis of diverse particles to maintain lung homeostasis and control potential infectious insults^3–5^. Cigarette smoking dramatically impairs AM phagocytic function^6,7^, as well as impacting the immunometabolic response to pathogens^8,9^, contributing to the increased susceptibility to infection seen in this population and in individuals with smoking-related chronic lung diseases^10^. With an estimated 1.25 billion tobacco users worldwide^11^ and growing concern over the health impacts of other types of particulate matters^12^, understanding the underlying pathobiology and exploring routes to correct this immune dysregulation is vitally important.

Murine AM largely originate from fetal liver monocytes, seeding to the lung after birth^13–15^, but are slowly replaced by circulating monocytes at an estimated rate of 40% per year^16^. Following LPS challenge, this rate of replacement increases to 85% over 2 months, findings supported by work in an influenza A-induced acute lung injury model^17^ and in a *Streptococcus pneumonia* infection model^18^. In humans, replenishment of the AM population by circulating monocytes has also been demonstrated^19^. Analysis of sex-mismatched lung transplant recipients showed that almost all donor AM derived from circulating recipient monocytes at one year post transplant^19^. Furthermore, children with hereditary CCR2 deficiency, responsible for monocyte trafficking to the lung, have approximately 50% fewer AM compared to healthy controls^20^. These studies highlight the importance of peripheral monocytes in replenishing the AM population and maintaining lung immunity. Interestingly, Corleis *et al*. observed increased proportions of airspace monocytes in the lungs of smokers compared to never-smokers, suggesting that cigarette smoke exposure may increase recruitment of circulating monocytes to the lung.

As precursors to the pivotal AM, these recruited airspace monocytes derived from circulating monocytes may present an attractive target for systemically administered host-directed therapy to enhance lung immunity. To harness this potential therapeutic target, however, an in-depth understanding of the biology, metabolism and function of these poorly characterised cells is required. Patel *et al*. previously developed a model of blood monocyte subset kinetics using human *in vivo* isotope labelling, wherein classical monocytes (CD14+CD16-) exit the bone marrow during inflammation, circulate for 1 day and then mostly leave the bloodstream^21^. A small population matures into intermediate monocytes (CD14+CD16+), which either leave the bloodstream or further develop into non-classical monocytes (CD14lowCD16+). More recently, Corleis *et al*. used RNA velocity analysis to identify a new monocyte marker, CD93, to identify a subset of newly recruited monocytes in the airspaces^22^. This marker of newly recruited airspace monocytes was rapidly lost during *in vitro* culture, limiting experimental characterisation of this niche population when removed from its *in vivo* environment.

The importance of cellular metabolism in supporting innate immune function is now widely appreciated^23^. Murine AM metabolism is well characterised and is dominated by PPAR-γ driven lipid metabolism pathways, which is linked to the lipid rich environment of the lungs^24–27^. This contrasts with murine peritoneal macrophages, where glucose metabolism is required for optimal phagocytic capacity^28,29^. In part due to the difficulty in studying tissue-resident immunology in humans, the metabolism of human AM is less well characterised, while that of airspace monocytes remains unexplored. It was recently shown that LPS or IFN-γ boosts human AM metabolism, particularly glycolysis^30^, and *in vitro Mtb* infection can alter both AM glycolysis and oxidative phosphorylation (OxPhos)^8,31–33^. Cigarette smoke dramatically alters baseline AM metabolism and downstream functions following infectious challenge^8,9,34^. Given the growing interest in immunometabolism as a target for host-directed therapies for the treatment of respiratory infections and other lung diseases^35,36^, enhanced understanding of human AM and airspace monocyte metabolism, and their vulnerability to cigarette smoke exposure, is urgently needed.

Here, we employ novel spectral flow cytometry-based metabolic and functional assays to show that *ex vivo* human AM and airspace monocytes have distinct metabolic phenotypes, with newly recruited CD93+ airspace monocytes exhibiting low dependency on mitochondrial metabolism and reduced phagocytic function. Boosting human monocyte OxPhos using the TCA cycle intermediate succinate enhanced phagocytosis, and crucially, drove their differentiation into macrophages with superior function. These findings highlight succinate as a therapeutic candidate for lung disease, with implications both in systemic monocyte-targeted therapies and metabolic preconditioning for adoptive cell therapies.

## Results

### CD93+ airspace monocytes represent an immature myeloid population in the human lung

To explore the immune landscape of the human lung, we developed multiparameter spectral flow cytometry panels for use on fresh *ex vivo* BAL samples. Due to the high autofluorescence of some lung resident populations^37^, we first explored the emission of the autofluorescence across the violet, blue, yellow/green and red channels of the Cytek Aurora flow cytometer. As expected, autofluorescence was high around the B2 channel (FITC channel), however we observed even higher levels of autofluorescence across the violet channels (Supp Fig. 1), while autofluorescence was lowest in the red channels. AM had the highest level of autofluorescence, followed by airspace monocytes and then lymphocytes (Supp Fig. 1A-B). Current smokers had higher levels of autofluorescence than ex-smokers (Supp Fig. 1C).

We designed a panel around the channels that were least impacted by autofluorescence, and used this to identify CD206+ AM, CD14+ monocytes, monocyte subsets and lymphocyte populations in the BAL samples (Supp Fig. 2A for gating strategy). Approximately 80% of total cells were identified as CD45+ leukocytes (Fig. 1A) and AM and granulocytes constituted 79.3% and 13.2% of the myeloid population respectively (Fig.1B). To explore myeloid recruitment to the lung, we examined the frequencies of airspace monocytes. As recently described^22^, we identified airspace monocytes as CD45+, FSC low, SSC low, CD14+ leukocytes (Supp Fig. 2A), and these represented 6.9% of the total myeloid cells (Fig. 1B). As expected, AM were bigger and more granular than both airspace monocytes and granulocytes (Supp Fig. 3A-B)^38^.

**Figure 1.**
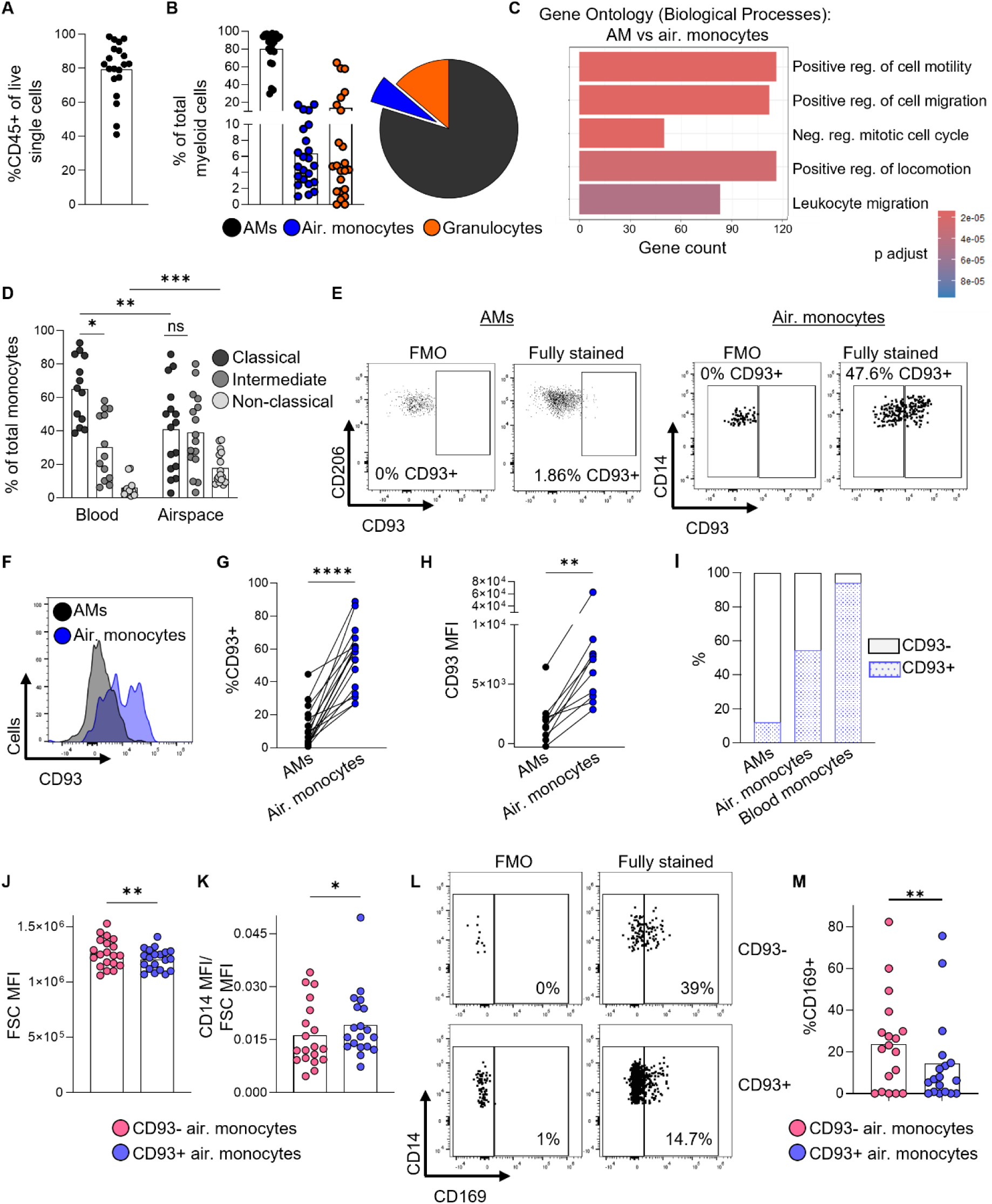
CD93+ airspace monocytes represent an immature myeloid population in the human lung. Fresh BAL samples were stained *ex vivo* and analysed by flow cytometry. (A) Frequency of CD45+ leukocytes from total live single cells. (B) Frequencies of AM, airspace monocytes and granulocytes from total myeloid cells. (C) Gene ontology analysis of scRNAseq data on fresh *ex vivo* BAL samples comparing AM vs airspace monocytes. (D) Frequencies of monocyte subsets in blood and BAL samples. (E-F) Representative dot plot and histogram of CD93 expression in AM and airspace monocytes. (G-H) Pooled data of the frequency of CD93 expression and CD93 MFI in AM and airspace monocytes. (I) Mean frequencies of CD93 expression in AM, airspace monocytes and blood monocytes. (J-K) FSC and CD14 MFI in CD93- and CD93+ airspace monocytes. (L-M) Representative dot plot and pooled data of the frequency of CD169 expression in CD93- and CD93+ airspace monocytes. (G-H and J-M) Data was compared using a paired t test. (D) Data was compared using a 2way ANOVA test. Air=airspace. N = 10-24, *p<0.05, **p<0.01, ***p<0.001, ****p<0.0001.

We investigated the biological processes that differ between AM and airspace monocytes using a publicly available single cell RNA sequencing dataset from fresh, *ex vivo* human BAL samples (n = 5 smokers and n = 4 never smokers)^22^. Gene ontology analysis of the differentially expressed genes highlighted several pathways involved in cell motility and migration (Fig. 1C), supporting the emerging concept that airspace monocytes have recently migrated from the blood to the lungs to replenish the resident AM population^18–21^. We explored the distribution of monocyte subsets within the total airspace monocyte population and compared it to that from healthy donor blood. In agreement with previous studies on human monocyte kinetics^21^, classical monocytes represented the greatest population in the blood, followed by intermediate and then non-classical monocytes (Fig. 1D). This contrasted with the distribution of monocyte subsets in the airspace, where we observed an expansion in intermediate and non-classical monocytes, and a concomitant decrease in the proportion of classical monocytes (Fig. 1D). As classical monocytes mature into intermediate monocytes and then non-classical monocytes^21^, this finding suggests that airspace monocytes are undergoing differentiation and maturation.

The cell surface adhesion receptor CD93 was recently described as marker of newly recruited monocytes in the airspaces^22^. The authors observed approximately 100%, 50%, and 0% CD93 expression in blood monocytes, airspace monocytes and AM respectively, and suggested that airspace monocytes downregulate CD93 during maturation in the lung. We sought to reproduce these findings in our patient cohort, and observed 94.1%, 54.3%%, and 12.1% CD93+ in blood monocytes, airspace monocytes and AM respectively (Fig. 1E-I). We also investigated CD93 expression in blood and airspace monocyte subsets and observed high CD93 expression in classical and intermediate monocytes, and lower CD93 expression in non-classical monocytes (Supp Fig. 3C). These data support the notion that CD93 expression is reduced during monocyte maturation. CD93+ airspace monocytes were smaller and expressed higher levels of CD14 compared to CD93- airspace monocytes (Fig. 1J-K), while the frequency of expression of the tissue residency marker CD169 was reduced (Fig. 1L-M). CD169 expression was also lower in classical airspace monocytes compared to the more mature intermediate and non-classical monocytes (Supp Fig. 3D). Together these findings suggest that CD93+ airspace monocytes are an immature monocyte population in the human lung and likely represent a newly recruited population.

### Smoking increases the frequency of airspace monocytes in the human lung

We next investigated biological and environmental factors that could impact recruitment of immune cells to the human lung. The cellular yield from BAL samples was similar between male and female donors and between donors <55 years old and ≥55 years old (Supp Fig. 3E-G). In contrast, BAL samples from smokers had significantly greater numbers of cells than those from never smokers, suggesting that smoking increases recruitment of immune cells to the lung (Fig. 2A-B). We next compared the myeloid populations from never smokers, ex-smokers and current smokers. While there were no significant differences in the frequencies of AM, smokers had increased frequency of airspace monocytes (Fig. 2C-D), and this was accompanied by an increase in the frequency of CD93+ airspace monocytes (Fig. 2E). Moreover, smoking impacted the distribution of monocyte subsets in the lung, with ex/current smokers containing more intermediate and non-classical monocytes than never smokers (Fig 2F).

**Figure 2.**
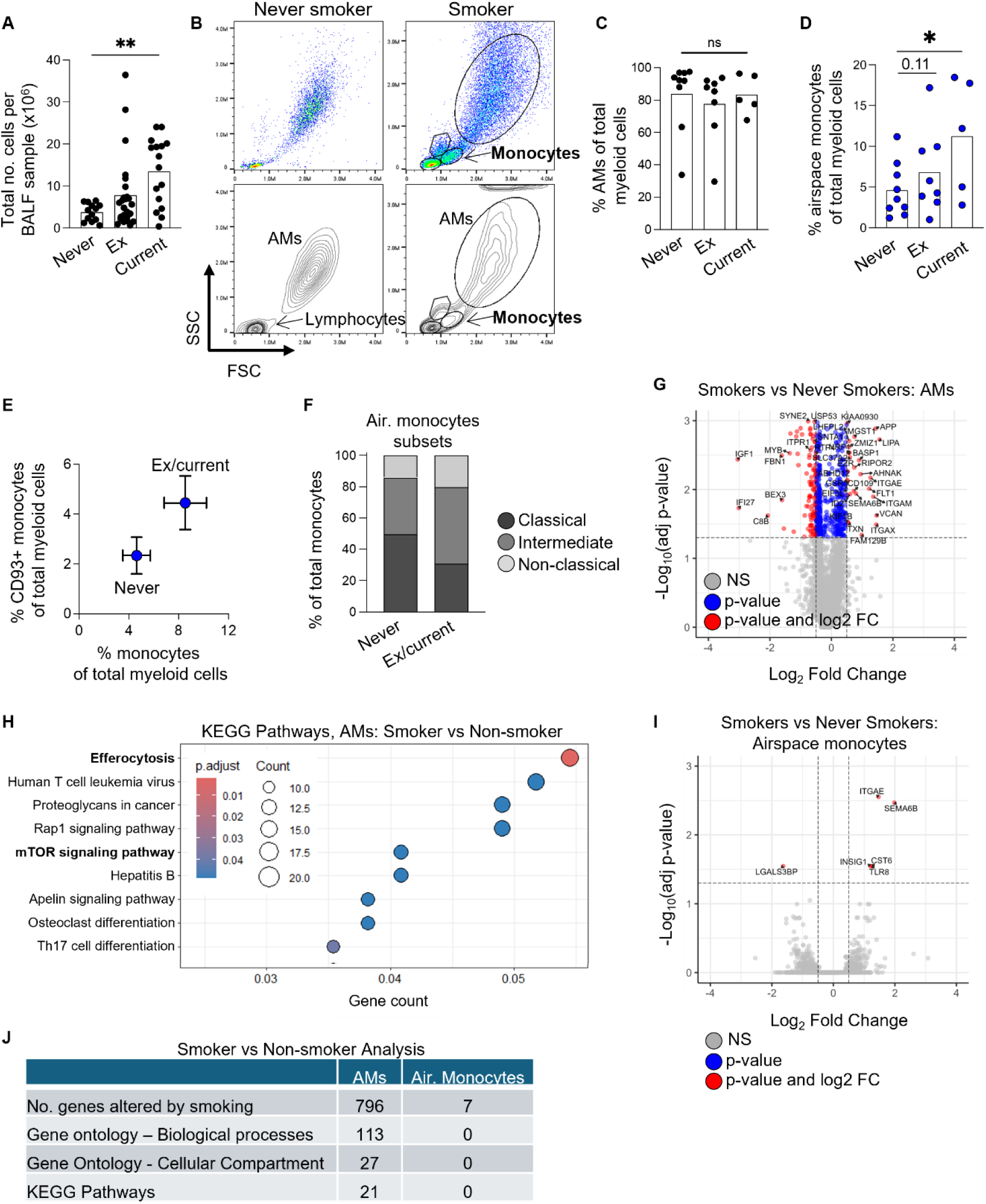
Smoking increases the frequency of airspace monocytes in the human lung. Fresh BAL samples were stained *ex vivo* and analysed by flow cytometry. (A) Total cellular yield of BAL samples according to smoking status. (B) Representative dot plot of leukocyte populations by FSC and SSC in a never smoker and smoker BAL sample. (C-E) Frequency of AM, airspace monocytes, and CD93+ monocytes of total myeloid cells according to smoking status. (F) Distribution of airspace monocyte subsets according to smoking status. (G) Differential gene expression analysis of scRNA sequencing data of *ex vivo* BAL samples between smokers and never-smokers in AM. (H) KEGG pathway analysis in AM between smokers and never-smokers. (I) Differential gene expression analysis of scRNA sequencing data of *ex vivo* BAL samples between smokers and never-smokers in airspace monocytes. (J) Analysis summary between smokers and never-smokers for AM and airspace monocytes. (A and C-D). Data was compared using a 2way ANOVA test. (F) Data was compared using an unpaired t test. ns = not significant. Air=airspace. N = 5-25, *p<0.05, **p<0.01.

We hypothesised that if airspace monocytes have recently migrated from the blood to the lung, they would be less impacted by smoking exposure than the more mature AM population. We analysed the Corleis *et al*. single cell RNA sequencing dataset^22^ and performed differential gene expression analysis on smokers versus never smokers. As expected, the transcriptome of AM was dramatically altered by smoking (Fig. 2G), and KEGG pathway analysis highlighted efferocytosis (phagocytosis of apoptotic cells) and mTOR signalling (important for metabolism) as key pathways impacted by smoking (Fig. 2H). In contrast, only 7 out of 14791 genes were altered by smoking in airspace monocytes, and there were no pathways significantly impacted by smoking when using gene ontology or KEGG analysis (Fig. 2I-J). Together these findings support the conclusion that airspace monocytes are a newly recruited population in the human lung and are enriched in response to smoking.

### CD93+ airspace monocytes have low mitochondrial dependency

Despite the fundamental role that metabolism plays in supporting immune functions in the lung^36^, little is known about the metabolism of human AM directly *ex vivo* and the metabolism of airspace monocytes is unexplored. We began our interrogation of AM and airspace monocyte metabolism by investigating the expression of key nutrient transporters using flow cytometry. AM had a higher frequency of Glut1 protein expression than airspace monocytes (Fig 3A). The mean expression level of Glut1 was also higher in AM compared to airspace monocytes (Supp Fig. 4A), however this was largely abrogated when Glut1 expression values were normalised to cell size, showing that high Glut1 expression in AM is linked to their large cell size (Fig. 3B). We examined the expression of CD98, a subunit of the solute carrier family 7 member 5 (SLC7A5) transporter which is important for uptake of large neutral amino acids such as arginine and tryptophan. Both AM and airspace monocytes had a high frequency of CD98 expression (Supp Fig. 4B), however the mean level of expression was highest in AM, and this was independent of cell size (Fig. 3C). We explored the expression of nutrient receptors in CD93- and CD93+ airspace monocytes. While expression of Glut1 was similar between the two populations (Supp Fig. 4C), CD93+ airspace monocytes had reduced expression of CD98 (Fig. 3E). These data suggest that CD98 expression increases as CD93+ airspace monocytes mature into CD93-airspace monocytes and then AM, implicating potential metabolic reprogramming during the monocyte to macrophages transition.

**Figure 3.**
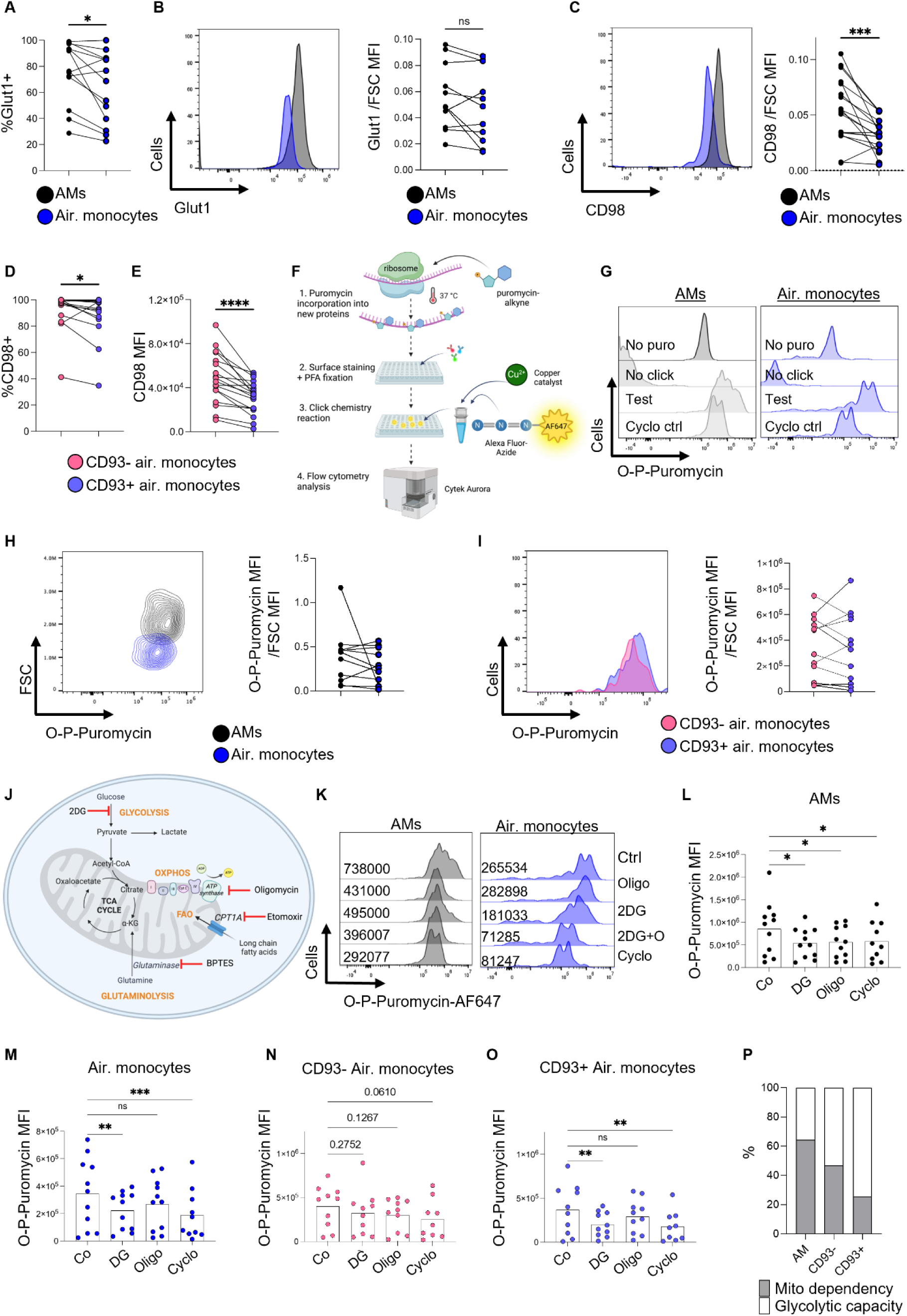
Divergent mitochondrial dependencies in human airspace monocytes and AM. Fresh BAL samples were stained *ex vivo* and analysed by flow cytometry. (A) Frequency of Glut1 expression and (B) representative histogram and pooled data for Glut1 MFI normalised to FSC MFI in AM and airspace monocytes. (C) Representative histogram and pooled data for CD98 MFI normalised to FSC MFI in AM and airspace monocytes. (D-E) CD98 expression in CD93- and CD93+ airspace monocytes. (F) Schematic of click chemistry-based *de novo* protein synthesis assay. (G) Representative histogram of O-P-Puromycin incorporation in AM and airspace monocytes, cycloheximide was used as a negative control. (H) Contour plot of O-P-Puromycin versus FSC, and pooled data for O-P-Puromycin MFI normalised to FSC MFI in AM and airspace monocytes. (I) O-P-Puromycin MFI normalised to FSC MFI in CD93- and CD93+ airspace monocytes. (J) Schematic of the targets of metabolic inhibitors used. *Ex vivo* cells were incubated with metabolic inhibitors (100mM 2DG, 1 µM oligo, 5 µM etomoxir or 5 µM BPTES) for 15 mins prior to *de novo* protein synthesis assay. (K) Representative histogram of O-P-Puromycin incorporation in AM and airspace monocytes with metabolic inhibitors. (L-O) Pooled data of O-P-Puromycin MFI in AM, airspace monocytes and blood monocytes. (P) Metabolic parameters calculated according to the SCENITH protocol. (A-E and H-I) Data was compared using a paired t test. (L-O) Data was compared using a 2way ANOVA test. Air=airspace. N = 10-19, *p<0.05, **p<0.01, ***p<0.001, ****p<0.0001.

To explore the metabolic dependencies of *ex vivo* AM and airspace monocytes at single cell level, we employed a modified version of SCENITH (Single Cell ENergetIc metabolism by profiling Translation inhibition)^39^. SCENITH takes advantage of the fact that over half of the ATP produced by a cell is immediately used for protein synthesis, meaning that substrates for ATP production can be interrogated by investigating the impact of metabolic inhibitors on *de novo* protein synthesis. *De novo* protein synthesis is measured by monitoring incorporation of puromycin into new protein chains^40^.

While SCENITH typically uses an anti-puromycin flow cytometry antibody for puromycin labelling, we optimised a click chemistry-based method to fluorescently label puromycin in AM and airspace monocytes, as this approach has reduced background staining which is necessary when working with the highly autofluorescent cells of the lung. In brief, cells were supplied with alkyne-bearing puromycin which was taken up and incorporated into new proteins. Cycloheximide, an inhibitor of protein translation, was used as a negative control. Following surface staining and PFA fixation, an azide-bearing fluorophore and copper catalyst were added, triggering the click chemistry reaction between the alkyne of the puromycin and the azide the of fluorophore, and the now fluorescently labelled puromycin was measured using flow cytometry (Fig. 3F-G).

We first compared the levels of *de novo* protein synthesis between AM and airspace monocytes *ex vivo*. While the frequency of puromycin incorporation was similar between the populations (Supp Fig. 4D-E), the mean levels of *de novo* protein synthesis was higher in AM than in airspace monocytes (Supp Fig. 4F-G). This difference could be attributed to their larger cell size, as normalising to cell size abrogated the significance of the difference (Fig. 3H). CD93- and CD93+ airspace monocytes had similar mean levels of *de novo* protein synthesis (Fig. 3I), however non-classical airspace monocytes had higher levels of *de novo* protein synthesis than classical airspace monocytes (Supp Fig. 4H).

We next investigated which nutrients are important for fuelling ATP production in these cells by analysing the impact of various metabolic inhibitors on the rates on *de novo* protein synthesis (Fig. 3J). In AM, treatment with 2-Deoxy-D-glucose (2DG) or oligomycin (oligo) inhibited protein synthesis to a similar extent as co-treatment with 2DG + oligo (Fig. 3K-L), suggesting a metabolic configuration wherein glucose-fuelled glycolysis and OxPhos are both essential for optimal ATP production and protein synthesis. In contrast, 2DG but not oligo inhibited protein synthesis in total airspace monocytes (Fig. 3K and M), suggesting that this population is primarily reliant on aerobic glycolysis to fuel ATP production and protein synthesis.

To gain insight into whether this metabolic dichotomy was due to newly recruited monocytes having unique metabolic dependencies, we carried out SCENITH analysis on CD93+ and CD93-airspace monocytes. The impact of metabolic inhibitors on CD93-airspace monocytes was variable, although there was a trend towards decreased protein synthesis with both 2DG and oligo (Fig. 3N). However, protein synthesis in CD93+ newly recruited airspace monocytes was highly sensitive to treatment with 2DG but not oligo (Fig. 4O). The metabolic profile of CD93+ newly recruited airspace monocytes mirrored that of blood monocytes, whereby only 2DG treatment significantly reduced protein synthesis (Supp Fig. 4I). Mitochondrial dependency was calculated which reflects sensitivity to oligo relative to 2DG+oligo^39^. AM had greater mitochondrial dependency than CD93-airspace monocytes, and these CD93-airspace monocytes had greater mitochondrial dependency than the more immature CD93+ airspace monocytes (Fig. 4P and Supp Fig. 4J).

**Figure 4:**
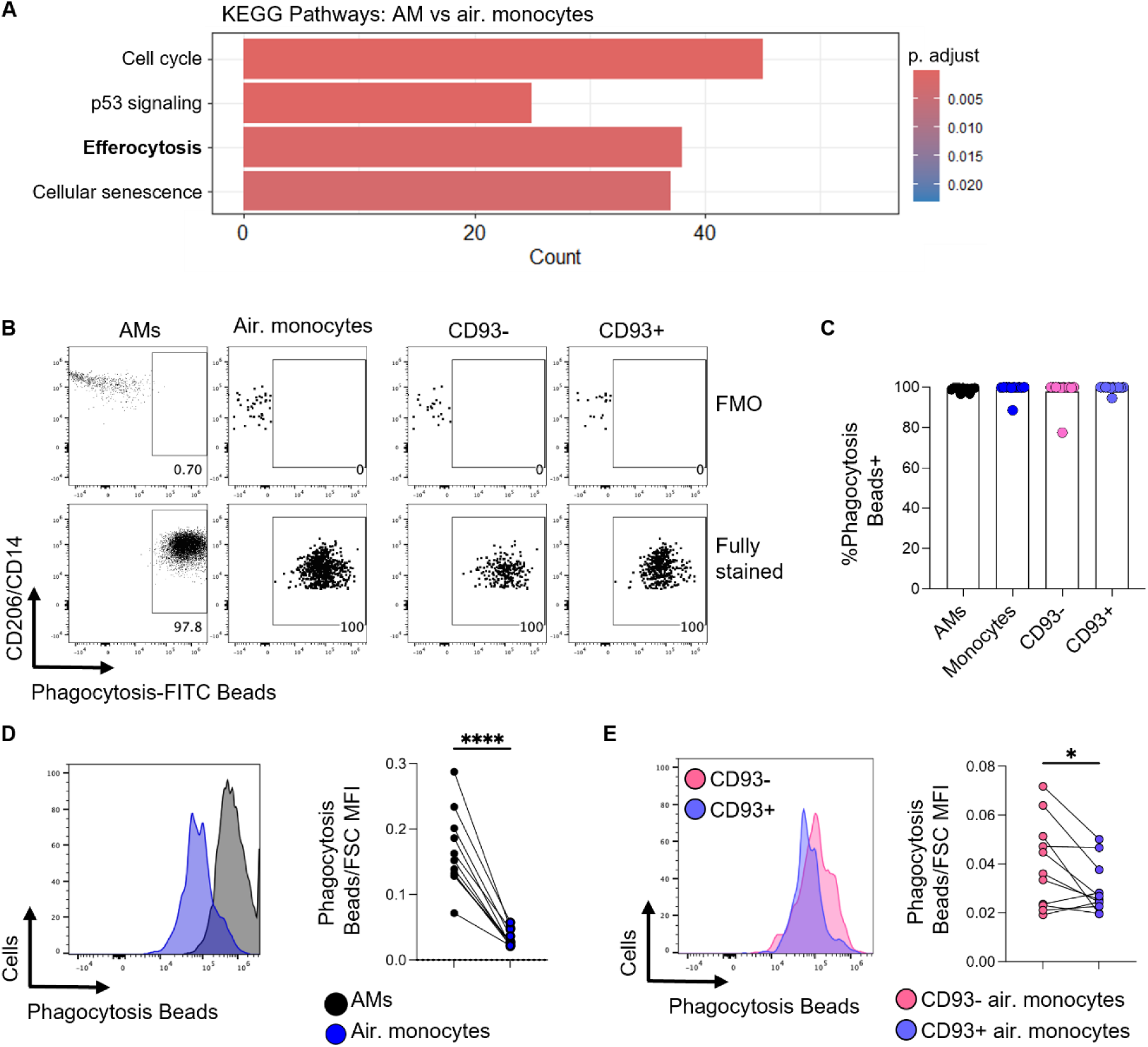
CD93+ airspace monocytes have low phagocytic capacity. (A) KEGG pathway analysis of scRNAseq data on fresh BAL samples between AM and airspace monocytes. (B-C) *Ex vivo* BAL cells were incubated with FITC-labelled phagocytosis beads for 50 mins and analysed by flow cytometry. Representative dot plot and pooled data of phagocytosis bead uptake in AM and airspace monocytes. (D) Representative histogram of phagocytosed FITC beads and pooled data of FITC bead MFI normalised to FSC MFI in AM and total airspace monocytes and in (E) CD93- and CD93+ airspace monocytes. (D-E) Data was compared using a paired t test. Air=airspace. N = 11, *p<0.05, ****p<0.0001.

Given the high mitochondrial dependence observed in AM, we next asked whether nutrients other than glucose contribute to this, as a variety of substrates can fuel mitochondrial OxPhos. To address this, we treated cells with low dose etomoxir (5μM) to inhibit the import of long chain fatty acids into the mitochondria through CPT1A, or with BPTES to inhibit glutamine anaplerosis of the TCA cycle (Fig. 3J). Interestingly, the impact of etomoxir and BPTES was highly variable in all populations analysed (Supp Fig. 4I-K), and SCENITH calculations did not show any difference in fatty acid oxidation/amino acid oxidation capacity between the groups (Supp Fig. 4L), suggesting that engagement in fatty acid oxidation or glutamine anaplerosis to fuel ATP production may be donor dependent, although this may be related to the short time in which the cells were exposed to the inhibitors (50 mins in total). Overall, these data highlight the unique metabolic profiles of *ex vivo* AM and airspace monocytes and suggest that immature airspace monocytes undergo metabolic reprograming to become more dependent on glucose-fuelled OxPhos as they mature and into AM.

### CD93+ airspace monocytes exhibit inherently low phagocytic capacity

We next asked whether the unique metabolic profile of airspace monocytes is associated with altered immune functions. Using the Corleis *et al.* dataset^22^, we carried out KEGG pathway analysis on the differentially expressed genes between AM and airspace monocytes and found that efferocytosis was highlighted a top pathway (Fig. 4A). To explore this further, we developed an assay wherein freshly isolated *ex vivo* BAL cells were cultured with FITC-labelled phagocytosis beads for 50 mins and then stained for surface markers and intracellular cytokines and analysed by flow cytometry. To start, we compared the phagocytic capacity of *ex vivo* AM and total airspace monocytes. Uptake of the phagocytosis beads was high for all populations, measuring ∼100% positive for FITC beads in each experiment (Fig. 4B-C). As expected, AM had a higher phagocytic capacity than airspace monocytes (Supp Fig. 5A), and this was also evident after normalisation to cell size (Fig. 4D). This suggests that the high mitochondrial dependence in AM may be associated with their high phagocytic capacity, and so we hypothesised that newly recruited CD93+ airspace monocytes, which have low mitochondrial dependence, might also have low phagocytic capacity. Indeed, CD93+ airspace monocytes had reduced phagocytic capacity compared to more mature CD93-airspace monocytes, and this was independent of cell size (Supp Fig. 5B and Fig. 4E).

AM from some donors expressed high levels of the pro-inflammatory cytokine IL1β *ex vivo*, and this was reduced in airspace monocytes (Supp Fig. 5C-E). Differences in TNFα and IL10 expression were variable between donors (Supp Fig. 5D) and there were no significant differences in intracellular cytokine levels between CD93+ and CD93-airspace monocytes (Supp Fig. 5F). Overall, these data highlight important differences in the immune functions of AM and airspace monocytes, particularly phagocytosis, which may be underpinned by their unique metabolic profiles.

### AM and CD93+ airspace monocytes engage in different metabolic pathways to fuel phagocytosis

To test whether the unique metabolic configurations of AM and airspace monocytes could be driving their different phagocytic capacities, we performed the *ex vivo* phagocytosis assay in the presence of metabolic inhibitors. 2DG and oligo both restricted phagocytosis in AM and CD93-airspace monocytes (Fig. 5A-B), indicating a reliance on both glycolysis and OxPhos for optimal phagocytosis. In contrast, only 2DG significantly inhibited phagocytosis in CD93+ newly recruited airspace monocytes (Fig. 5C), a pattern also observed in blood monocytes (Fig. 5D), suggesting these populations depend primarily on glycolysis to fuel phagocytosis.

**Figure 5.**
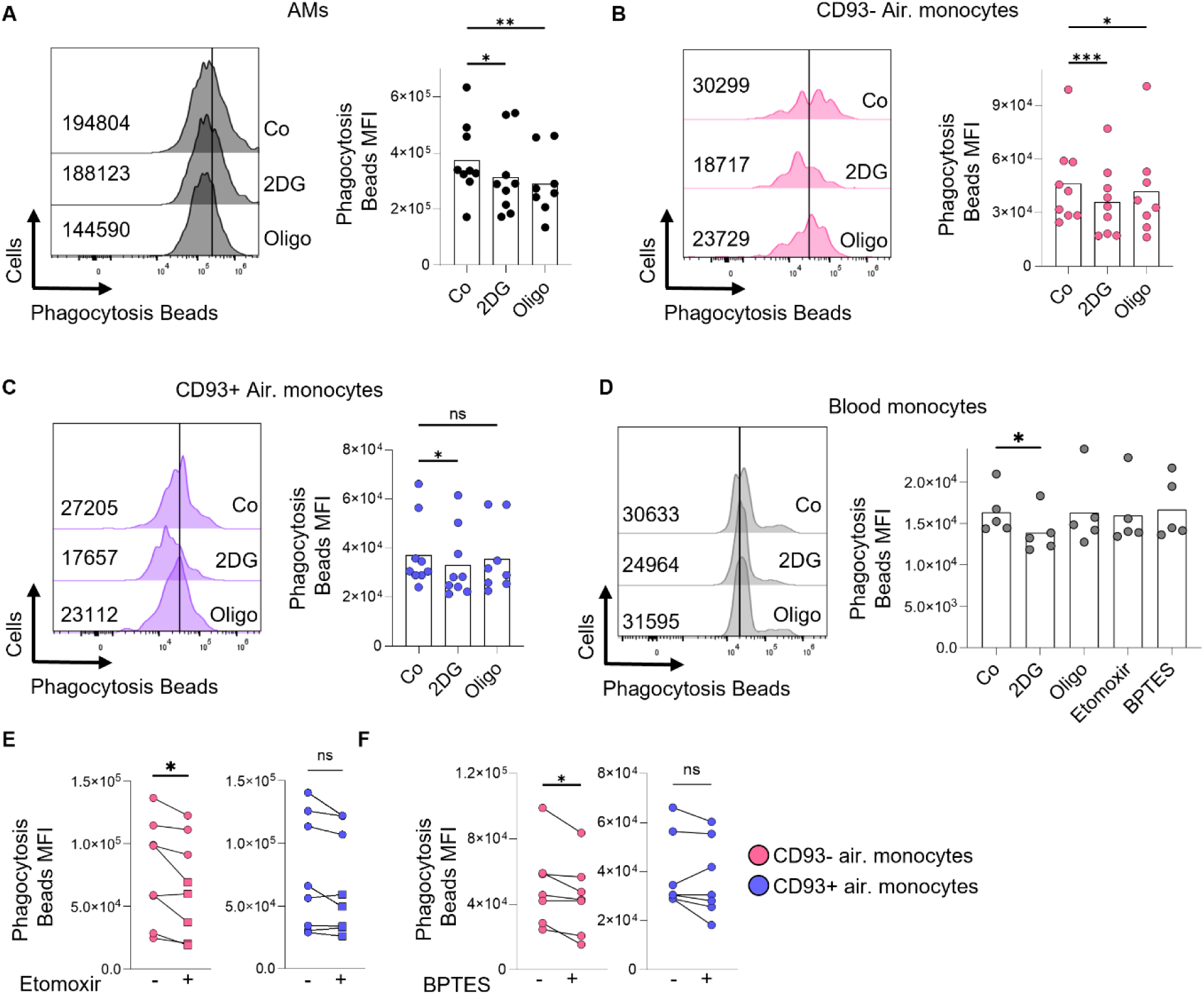
AM and CD93+ airspace monocytes engage in different metabolic pathways to fuel phagocytosis. *Ex vivo* cells were incubated with FITC-labelled phagocytosis beads for 50 mins in the presence or absence of metabolic inhibitors and analysed by flow cytometry. (A-C) Representative histogram and pooled data for AM and airspace monocyte phagocytosis ± 2DG (100 mM) or oligo (1 µM). (D) Representative histogram and pooled data for blood monocyte phagocytosis ± inhibitors. (E-F) Pooled data for CD93- and CD93+ airspace monocyte phagocytosis ± etomoxir (5 µM) or BPTES (5 µM). (A-D) Data was compared using a 2way ANOVA test. (E-F) Data was compared using a paired t test. Air=airspace, ns = not significant. N = 5-9, *p<0.05, **p<0.01, ***p<0.001.

Given that CD93-airspace monocytes relied on OxPhos for phagocytosis, we tested whether mitochondrial substrates like fatty acids or glutamine were required. Inhibition of CPT1A and glutaminase, key enzymes for fatty acid oxidation and glutamine anaplerosis respectively, diminished phagocytosis in CD93-airspace monocytes but not CD93+ airspace or blood monocytes (Fig. 5D-F). Altogether, these findings demonstrate that distinct immune cell populations in the lung employ different metabolic programme to support phagocytosis; CD93-airspace monocytes uniquely rely on OxPhos fuelled by glucose, fatty acids, and glutamine, alongside glycolysis, while the lack of mitochondrial engagement during phagocytosis in CD93+ airspace monocytes likely underlies their limited phagocytic capacity.

### Fuelling mitochondrial metabolism boosts phagocytic capacity in immature monocyte populations

AM and CD93-airspace monocytes rely on mitochondrial metabolism, which is linked to heightened phagocytic capacity compared to CD93+ airspace monocytes. We therefore hypothesised that boosting mitochondrial metabolism could enhance phagocytosis in immature blood monocytes and CD93+ airspace monocytes. Succinate plays a central role in the TCA cycle where it is oxidised to fumarate by succinate dehydrogenase (complex II of the electron transport chain), donating two electrons to ubiquinone^41^. Through this mechanism, succinate supports mitochondrial metabolism by simultaneously driving OxPhos and sustaining TCA cycle activity (Fig. 6A). We first tested whether succinate could boost OxPhos in human blood monocytes using seahorse metabolic flux analysis. Monocytes were plated directly *ex vivo*, media or succinate was injected into the well, and oxygen consumption was measured in real time. Succinate injection induced a small but consistent acute increase in oxygen consumption, indicating that human monocytes are poised to harness succinate as a mitochondrial substrate (Fig. 6B-C). The subsequent mito stress test demonstrated that monocytes exposed to succinate had greater maximal respiratory capacity and spare respiratory capacity (Fig. 6D-E), highlighting an enhanced ability to respond to increased energy demand in the presence of succinate. We next tested the impact of this enhanced mitochondrial metabolism on the monocytes’ phagocytic capacity. We repeated the *in vitro* phagocytosis assay and supplemented the media with succinate. Succinate supplementation boosted phagocytosis in both blood and CD93+ airspace monocytes (Fig. 7F-H), providing proof of concept that mitochondrial metabolism can be targeted to modulate phagocytosis in immature human monocyte populations.

**Fig 6:**
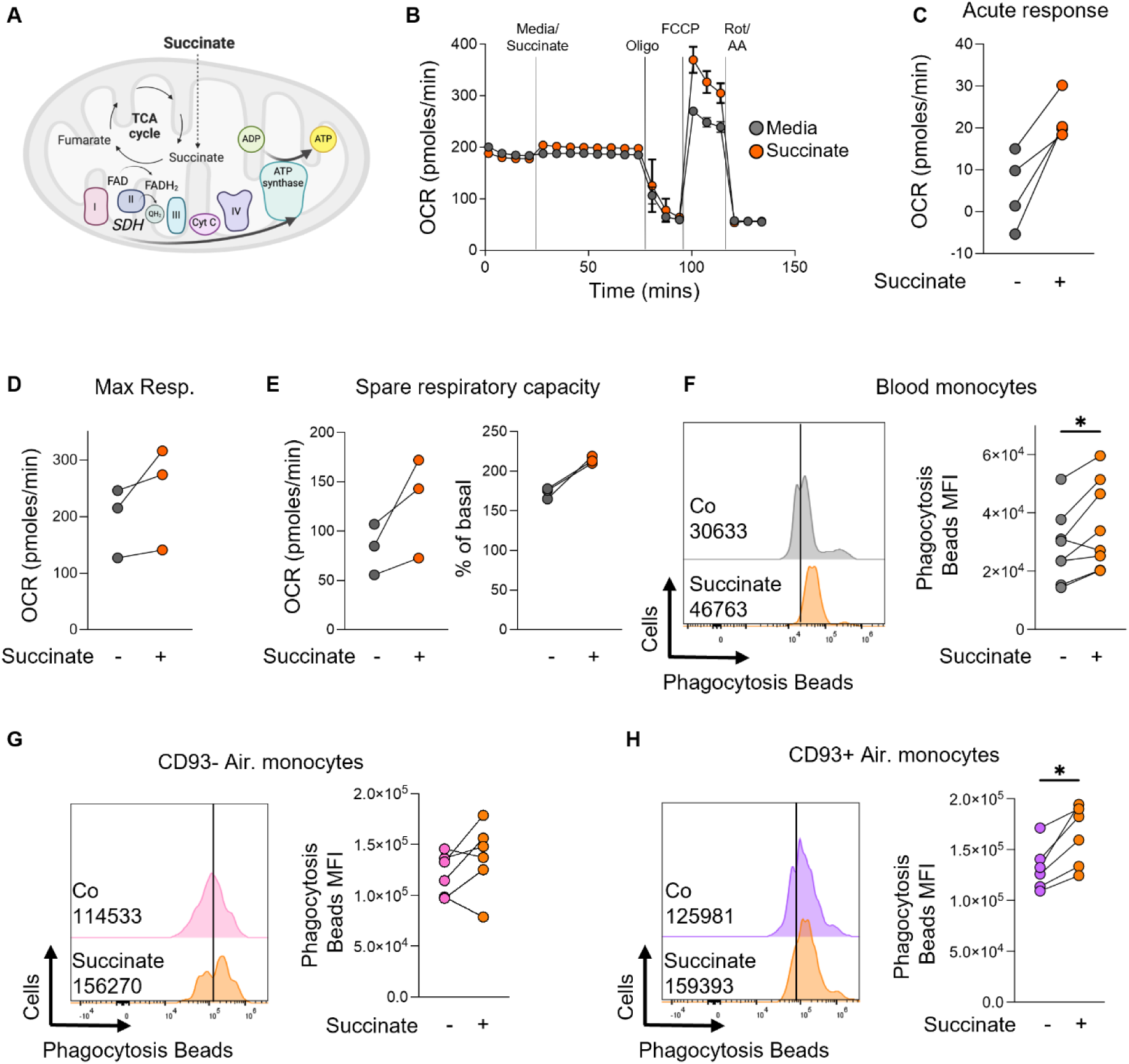
Fuelling mitochondrial metabolism boosts phagocytosis in blood and CD93+ airspace monocytes. (A) Schematic showing succinate fuelling of mitochondrial metabolism. (B) Seahorse metabolic flux analysis of OCR in *ex vivo* purified blood monocytes injected with seahorse media or succinate (5 mM), followed by oligo (2 µM), FCCP (0.5 µM) and rotenone + antimycin A (0.5 µM + 0.5 µM), measured in an XFp Analyzer. (C-E) Seahorse parameters calculated using Seahorse analytics. (F-H) *Ex vivo* blood or airspace monocytes were incubated with FITC-labelled phagocytosis beads for 50 mins in the presence or absence of succinate (5 mM), stained and analysed by flow cytometry. (F-H) Data was compared using a paired t test. SDH = succinate dehydrogenase, OCR = oxygen consumption, Air=airspace. N = 3-8, *p<0.05, **p<0.01, ***p<0.001.

**Fig 7:**
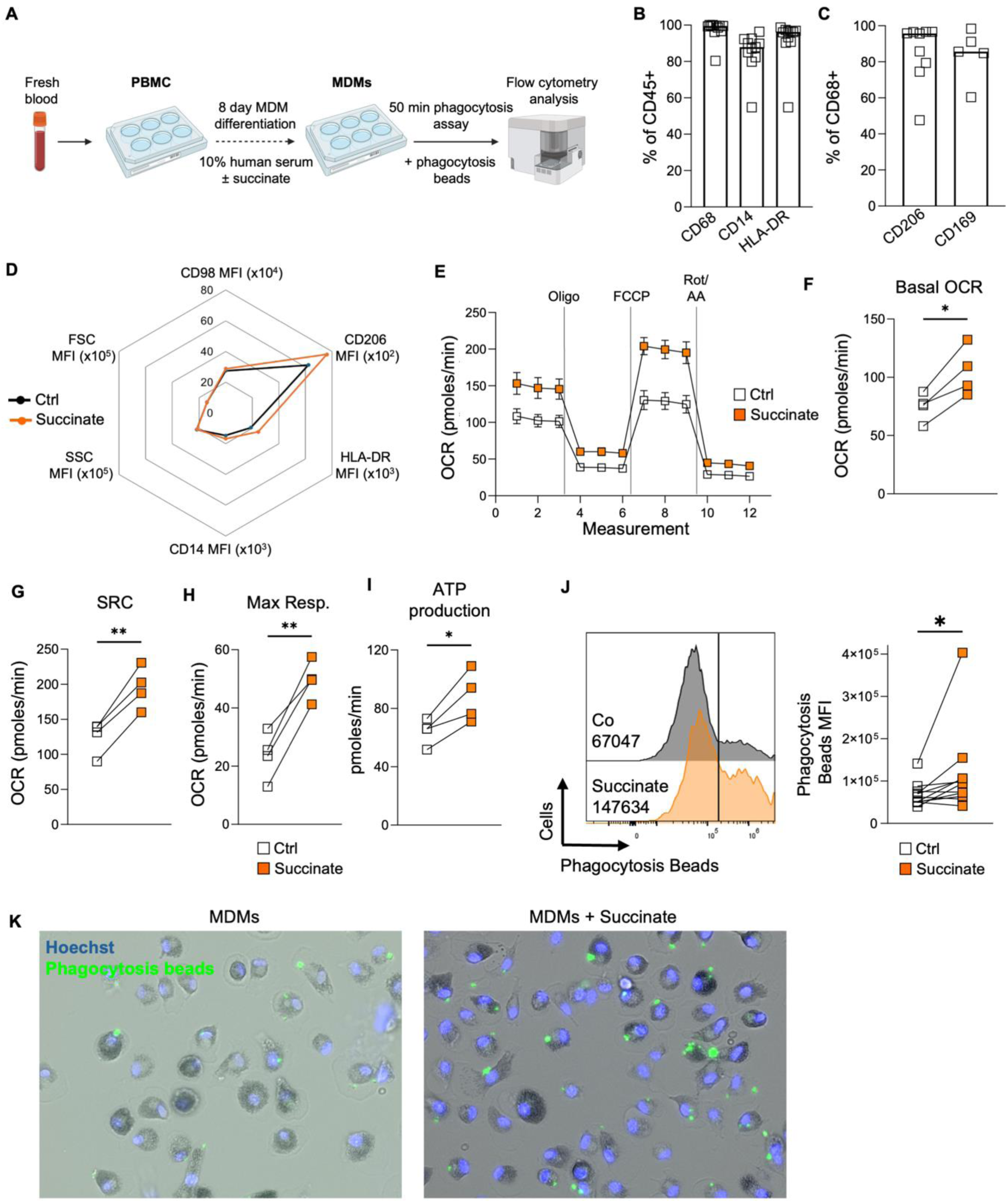
Succinate drives the differentiation of macrophages with enhanced metabolism and function. (A) Schematic showing *in vitro* MDM expansion protocol. (B-C) MDMs were stained and analysed by flow cytometry. (D) MDMs were expanded in the presence of absence of succinate (5 mM), stained and analysed by flow cytometry. Dots show the mean MFI expression value. (E) Seahorse metabolic flux analysis of OCR with injections of oligo (2 µM), FCCP (0.5 µM) and rotenone + antimycin A (0.5 µM + 0.5 µM), measured in an XFe96 Analyzer. (F-I) Seahorse parameters calculated using Seahorse analytics. (J) Representative histogram and pooled data of phagocytosis in MDMs expanded in the presence of absence of succinate. (K) Representative images of phagocytosis in MDMs stained with Hoechst and imaged using the Lionheart FX microscope. (F-J) Data was compared using a paired t test. MDM = monocyte-derived macrophage, SRC = spare respiratory capacity, OCR = oxygen consumption. N=4-11, *p<0.05, **p<0.01.

### Succinate drives the differentiation of macrophages with enhanced phagocytic function

Given that a central role of monocytes is to differentiate into macrophages upon recruitment into tissues, we investigated the impact of succinate supplementation on this process. PBMC were isolated from fresh blood and cultured in 10% human serum with or without succinate for 8 days (Figure 7A). The resulting monocyte-derived macrophages (MDMs) were then analysed by flow cytometry, revealing a highly pure population of MDMs that expressed of high levels of the canonical macrophage markers CD68, CD14 and HLA-DR (Figure 7B). CD68 was used to define the MDM population (Supp Fig. 5F), within which we also observed high expression of the maturity markers CD206 and CD169 (Figure 7C). Succinate supplementation did not significantly impact the phenotype of the MDMs, as evidenced by unchanged expression of HLA-DR and CD14 and unaffected cell size and granularity (Figure 7D). However, there was a trend towards increased expression of CD206, suggesting that succinate enhances MDM maturation (Fig. 7D and Supp Fig. 5G).

We investigated the metabolism of MDMs differentiated in succinate using seahorse flux analysis. Indeed, expansion of MDMs in the presence of succinate resulted in heightened mitochondrial metabolism, with basal metabolism, spare respiratory capacity and maximal respiration boosted in succinate-expanded MDMs compared to controls (Fig. 7E-H). This was also associated with increase ATP production (Fig. 7I) and shows that succinate exposure drives the development of highly oxidative MDMs. Finally, we carried out the phagocytosis assay and found that succinate supplementation produced MDMs with enhanced phagocytic capacity (Fig. 7J), and this was also confirmed using microscopy (Fig. 7K). Overall, these findings highlight monocyte mitochondrial metabolism as a key therapeutic target for enhancing monocyte and macrophage metabolism and phagocytosis and identify succinate as a candidate for boosting myeloid immunity in the blood and lungs.

## Discussion

The AM is the sentinel innate immune cell in the human lung, guarding this physiologic gateway against a constant insult of pathogens and pollutants, while maintaining environmental homeostasis within the airspace. Smoking dramatically impairs AM function, and smokers are vulnerable to a range of infectious and non-infectious diseases related to this dysfunction. Peripheral monocytes replenish and maintain the resident AM population^16,18–20,42^, playing a central role in supporting lung immunity, however their basic biology and metabolic regulation remains poorly understood. While immunometabolism has emerged in recent years as a central plank in innate immune function, metabolic interrogation of human tissue-resident cells such as the AM and airspace monocyte is limited by the profound impact *in vitro* culture exerts on cellular metabolism. Here, we provide an in-depth characterisation of the function and metabolism of freshly isolated *ex vivo* human airspace monocytes and AM, optimising cutting edge flow cytometry-based techniques to facilitate interrogation. We identify OxPhos as a critical pathway governing phagocytic capacity of lung myeloid cells. Acute targeting of mitochondrial metabolism using succinate boosted phagocytosis in both blood and airspace monocytes, highlighting monocyte-targeted metabolic therapies as a potential strategy for modulating immunity in the lung.

Through this work, we identify unique metabolic profiles of blood and airspace monocytes and AM. While blood monocytes and CD93+ airspace monocytes had low mitochondrial dependency, more mature CD93-airspace monocytes and AM were dependent on mitochondrial metabolism for both protein synthesis and phagocytosis. These observations align with those by Ehlers *et al*., who showed that human neonatal monocytes rely on OxPhos during differentiation, whereas monocytes from older individuals shift toward glycolysis^43^. Similarly, Wohnhaas *et al*. found that monocyte-derived AM in mice exhibit reduced expression of mitochondrial metabolic genes compared to resident AM^44^. This metabolic shift likely reflects the differing demands of each population. Glycolysis supports rapid anabolic biosynthesis and biomass expansion which is needed during monocyte differentiation. Once matured, macrophages benefit from mitochondrial metabolism’s higher ATP yield and ability to use diverse fuel sources like fatty acids and amino acids. Mitochondrial metabolism has also been shown to support efferocytosis^45,46^ as well as T cell longevity^47^, and may similarly contribute to the extended lifespan of AM. Overall, our results suggest that immature airspace monocytes undergo metabolic reprogramming to become increasingly reliant on mitochondrial metabolism as they mature into resident AM.

Consistent with Corleis *et al*.^22^, we found increased frequencies of airspace monocytes in smokers, with a similar trend in ex-smokers. This aligns with murine data showing elevated monocyte-derived AM following smoke exposure^44^. Chronic smoke-induced inflammation likely drives monocyte recruitment^48^, and we speculate that this may also occur in other chronic inflammatory lung diseases such as cancer, COPD and cystic fibrosis. Whether the increased proportion of airspace monocytes solely reflects increased recruitment, or whether these monocytes are arrested in their differentiation leading to accumulation is not clear – however, it is well established that smokers’ BAL fluid yields increased cell counts^49^, supporting the hypothesis that recruited monocytes are differentiating into mature AM. Furthermore, by boosting mitochondrial metabolism through succinate supplementation, we succeeded in driving blood and airspace CD93+ monocytes towards more functional similarity to mature CD93-monocytes and AM, showing that these an may be metabolically manipulated to promote differentiation towards a mature functional AM.

Interestingly, transcriptomic analysis of airspace monocytes at baseline revealed that smoking altered only 7 genes, in contrast to 796 genes altered by smoking in the AM transcriptome. The impact of smoking on AM function is well established^50^, and has been linked to cellular oxidative stress^9^. We previously showed that compared to non-smokers, smokers’ AM have reduced OxPhos *ex vivo*^8^ as well as diminished bioenergetic and cytokine response following *Mtb* infection^8,49^. Smoke-induced transcriptomic alterations have been reported in macrophages derived from murine bone marrow precursors in smoke-exposed mice^51^, as well as functional differences in monocyte-derived macrophages in human smokers^52^, suggesting that cigarette smoke can exert effects on innate immunity beyond the pulmonary compartment. However, our data suggest that recruited monocytes may be more protected from the harmful effects of cigarette smoke than the mature tissue-resident AM, and could thus offer an opportunity for pre-differentiation therapeutic intervention to enhance metabolism and function of the myeloid compartment in the smokers’ lung.

Given that our work highlights a critical link between mitochondrial dependency and phagocytic capacity in AM and airspace monocytes, we explored strategies to boost mitochondrial metabolism in immature blood and CD93+ airspace monocytes, to exploit this potential therapeutic target. Succinate is a key TCA cycle intermediate in macrophages, participating in metabolic reactions and modulating inflammatory signalling pathways^41,53,54^, however its acute impact on *ex vivo* human monocytes is unexplored. We demonstrate that succinate supplementation enhances mitochondrial metabolism and concurrently boosts phagocytosis in both blood and airspace monocytes. These findings align with research by Giorgi-Coll *et al*. showing that succinate improves OxPhos in metabolically stressed human microglia^55^, and by Ryan *et al*. who reported that NRF2 activation restores TCA cycle function and efferocytosis in dysfunctional AM from COPD patients^56^.

Remarkably, exposure of monocytes to succinate drove their differentiation into macrophages with superior phagocytic capacity, suggesting that systemic delivery of monocyte-targeted metabolic therapies could support the seeding of AM with improved functional and disease-preventive properties. This is particularly relevant in the context of smoking where AM have reduced OxPhos^8^. Regarding the practicality of using succinate as a treatment, Jung *et al.* recently showed in mice that oral administration of succinate is safe and results in its distribution throughout the blood and various tissues including the heart, liver, adipose and brain, which could still be detected after 4 hours^57^. In humans, delivery of succinate directly to the brain improved metabolic function in patients with traumatic brain injury, particularly those with mitochondrial dysfunction^58^. Moreover, Lu *et al.* recently demonstrated that succinate can be encapsulated within tumour-derived microparticles, which, when administered *in vivo*, promote M1 macrophage polarisation and effectively treat pre-clinical cancer models^59^. Thus, while there is a myriad of strategies to promote mitochondrial metabolism^60^, succinate itself is emerging as a potential candidate therapeutic agent and could provide a safe and low-cost approach to boosting lung immunity.

Beyond the concept of targeting monocytes systemically, our work has further implications for adoptive cellular therapies, particularly chimeric antigen receptor (CAR) macrophages which are undergoing intense investigation for the treatment of both haematological and solid tumour cancers, largely due to their potent phagocytic activity and favourable safety profile^61–64^. Succinate could potentially be leveraged to enhance the expansion of CAR macrophages with improved mitochondrial metabolism and phagocytic function, thereby increasing their therapeutic efficacy.

Overall, our study provides a comprehensive characterisation of human airspace monocytes and AM, uncovering mitochondrial metabolism as a key regulator of phagocytic function. By demonstrating that acute metabolic intervention with succinate enhances both monocyte mitochondrial activity and phagocytosis, we offer new insight into how immune function in the lung can be modulated. These findings may the pave the way towards innovative therapeutic strategies for treating lung disease, from systemic monocyte-targeted therapies to support AM function, to metabolic pre-conditioning of CAR-macrophages for adoptive cell therapy.

## Methods

### Cell culture

Peripheral blood mononuclear cells (PBMC) were isolated by density-gradient centrifugation over Lymphoprep (StemCell Technologies). Cells were resuspended in RPMI (Gibco) supplemented with 10% FBS (Gibco) and pen/strep.

All bronchoalveolar lavage (BAL) fluid donors were undergoing clinically indicated bronchoscopy and consented for additional BAL fluid to be acquired during their procedures (n = 32 patients). Patient demographic data, relevant medical diagnoses, medications, and smoking status were recorded (see **Table 1** and **Supp Fig. 6** for patient information). Exclusion criteria included lung cancer, HIV or HCV infection and active respiratory infection. The protocol for acquisition has previously been reported^65^. BAL fluid was filtered through a 100µm nylon cell strainer (Fisher) and centrifuged at 290g for 15 min. The pellet was resuspended in 2 ml RPMI (Gibco), supplemented with 10% FBS (Gibco), fungizone (2.5 μg/ml; Gibco) and cefotaxime (50 μg/ml; Melford Biolaboratories). Where indicated, 100 mM 2DG, 1 µM oligomycin, 5 µM Etomoxir, 5 µM BPTES, 100 µg/ml cycloheximide, 5 mM succinate or 6 mM glutamine was added.

**Table 1.** Patient information for BAL samples.

|  |  |
| --- | --- |
| **Total subjects** | n = 32 |
| **Sex distribution** | n = 19 Female, n = 13 Male |
| **Age range** | 23–84 years |
| **Mean age (years)** | 58.69, StDev = 13.8 |
| **Smoking status** | Never: n = 12 |
|  | Ex-smoker: n = 12 |
|  | Current smoker: n = 5 |
|  | Unknown: n = 4 |
| **Common respiratory diagnoses** | COPD: n = 5 |
|  | ILD: n = 6 |
|  | Sarcoidosis: n = 4 |
|  | Asthma: n = 3 |
| **Other diagnoses** | Atelectasis, pneumonia, haemoptysis, lung nodules, sinus disease, IPF, scleroderma, Wegener's granulomatosis, nasal polyp |

StDev = standard deviation, COPD = chronic obstructive pulmonary disease, ILD = interstitial lung disease, IPF = idiopathic pulmonary fibrosis.

### MDM differentiation protocol

Peripheral blood mononuclear cells (PBMC) were isolated from the blood of healthy donors by density gradient centrifugation and seeded at 2.5×10^6^ cells/mL on non-treated tissue culture plates (Costar). Over an 8-day differentiation, monocyte derived macrophages (MDM) were cultivated via plastic adherence in RPMI 1640 (Gibco) with 10% AB human serum (Sigma). To lift adherent MDM on day 8, the media was replaced with 1mL of ice-cold DPBS (Sigma) and incubated at 4°C for 30 minutes.

### Flow cytometry

Cells were seeded into V bottomed 96 well plates, Fc blocked (1/50, BioLegend) and stained for viability (Zombie NIR™ Fixable Viability Kit, BioLegend, 1/500) in PBS on ice for 12 mins. Cells were stained for surface markers (Table 2) for 20 mins on ice in FACS buffer (PBS + 2% FCS). Cells were fixed (IC Fixation Buffer, Invitrogen) for 20 mins on ice. Cells were stained for intracellular markers (Table 2) overnight at 4°C in permeabilization buffer (eBioscience). Cells were washed and resuspended in 200 μl FACS buffer. Samples were acquired on an Aurora flow cytometer (Cytek Biosciences).

**Table 2.**
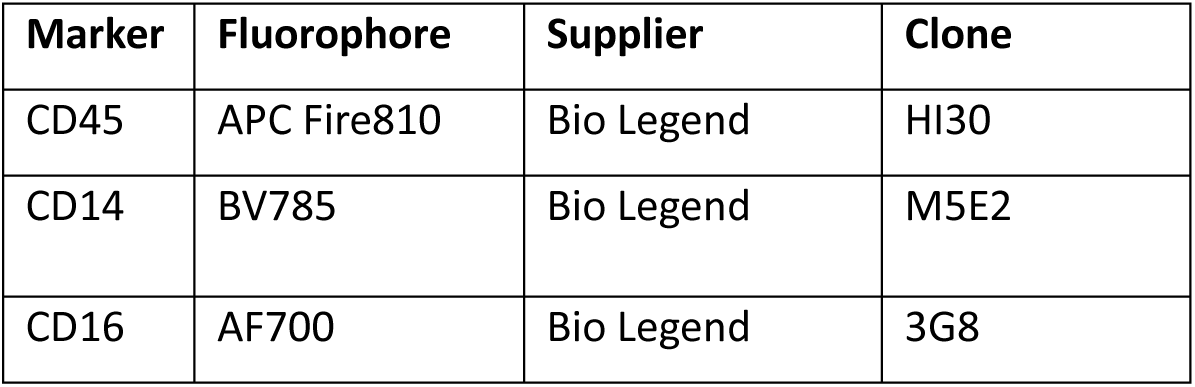
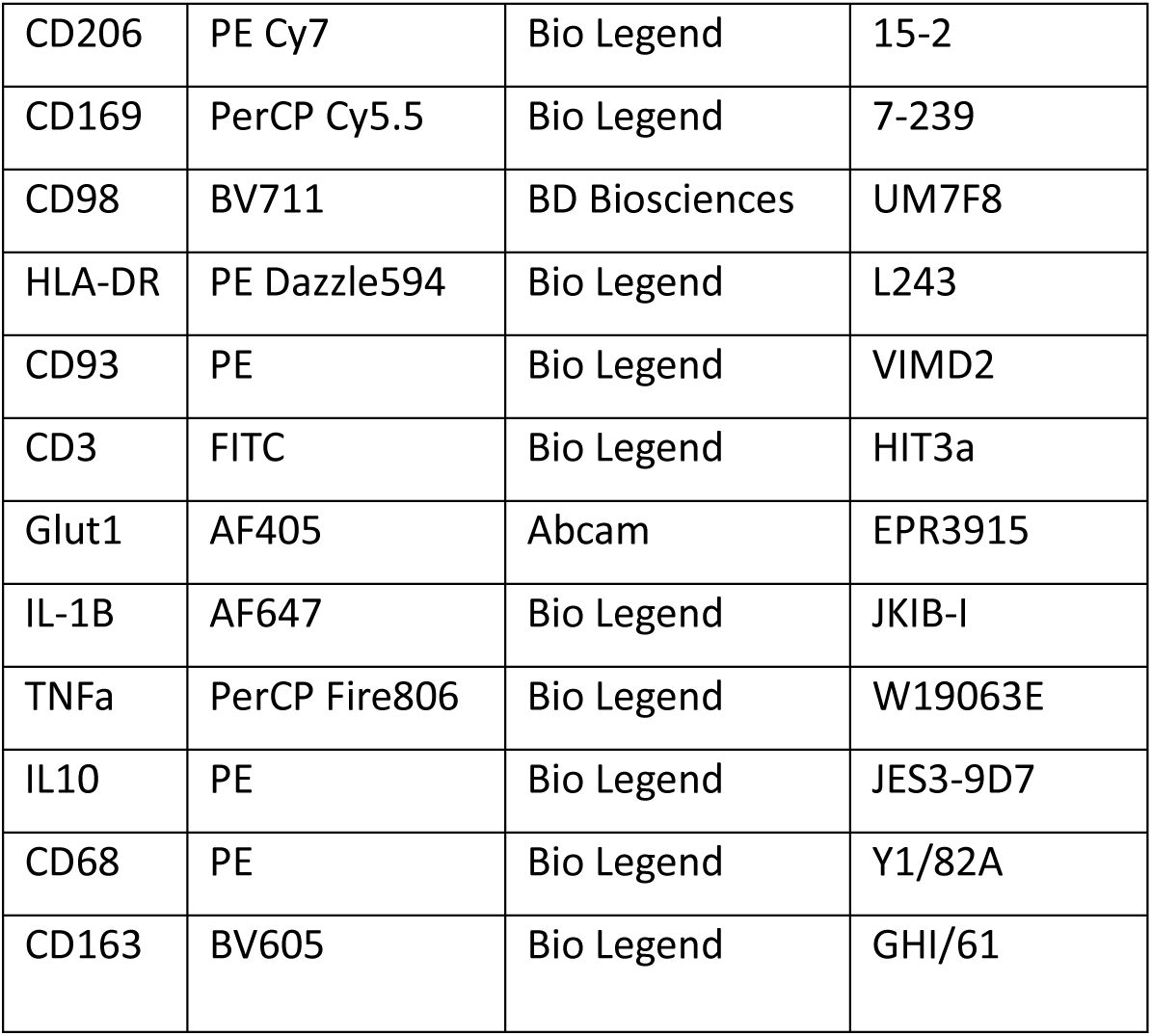
Flow cytometry antibodies.

### Single cell RNA sequencing analysis

Raw counts, metadata and cell type annotations were obtained from^22^, and loaded into the Seurat V5 R package^66^ for pseudobulk analysis. The Seurat FindMarkers() function was used to aggregate, for each donor, the raw counts of all cells of the same type. Then, the Seurat FindMarkers() function was used for differential expression analysis, with the argument test.use=DESeq2. False Discovery Rate was controlled using the Benjamini-Hochberg method, with Q = 0.05. The statistically significant differentially expressed genes from each comparison were used for downstream analysis. Gene Ontology enrichment analysis was performed using the clusterProfiler R package^67^. The enrichGO() and enrichKEGG() functions were used for Gene Ontology and KEGG Pathway enrichment analysis, respectively.

### Click chemistry-based *de novo* protein synthesis assay

Cells were seeded into 96 well plates in 37°C Hanks’ Balanced Salt Solution and pre-warmed alkyne-puromycin (O-propargyl-puromycin, OPP) was added (20 µM, Jena Bioscience). Cells were incubated for 30 mins at 37°C, washed and stained for flow analysis as described above. After the intracellular stain, cells were washed twice in PBS and the click chemistry reaction was performed. The click mix was made up in PBS by adding the following reagents in this exact order; 1 mM copper (II) sulfate pentahydrate, 10 mM sodium l-ascorbate, 1 mM tris-hydroxypropyltriazolylmethylamine (THPTA), 160 mM aminoguanidine hydrochloride, and 2.5 µM AZDye 647 Azide. Cells were kept at room temperature in the dark for 1 hour, washed twice in PBS and resuspended in FACS buffer. Samples were acquired on an Aurora flow cytometer (Cytek Biosciences).

### SCENITH

Working stocks for inhibitors were made in HBSS and pre-warmed to 37°C. After cells were seeded, pre-warmed metabolic inhibitors were added; 100mM 2DG, 1 µM oligomycin, 100mM 2DG + 1 µM oligomycin, 5 µM Etomoxir, 5 µM BPTES or 100 µg/ml cycloheximide. Cells were incubated for 15 mins at 37°C. After the 15 mins, OPP was added, and cells were incubated for 30 mins at 37°C. *De novo* protein synthesis was measured as described above. Metabolic parameters were calculated as previously described, according to following formulas:

-Glucose dependence = 100*((Co – 2DG)/(Co-DGO)).

-Fatty acid and amino acid oxidation capacity = 100 - glucose dependence.

-Mitochondrial dependence = 100*((Co – Oligo)/(Co-DGO)).

-Glycolytic capacity = 100 - mitochondrial dependence.

### Phagocytosis assay

Cells were seeded into 96 well plate in warm RPMI (Gibco), supplemented with 10% FBS (Gibco), fungizone (2.5 μg/ml; Gibco) and cefotaxime (50 μg/ml; Melford Biolaboratories) and the phagocytosis assay was performed according to the manufacturer’s instructions (Cambridge Bioscience). In brief, FITC-labelled latex beads were added (1/400) and cells were incubated for 50 mins at 37°C. For metabolic analysis, the following inhibitors were added for the duration of the assay: 100mM 2DG, 1 µM oligomycin, 5 µM Etomoxir or 5 µM BPTES. Cells were washed and stained for flow cytometry analysis as described above.

### Seahorse Analysis

#### Monocytes

For ex vivo monocyte seahorse analysis, monocytes were isolated from fresh blood using the Easysep human monocyte negative selection kit (StemCell). 0.2×10^6^ monocytes were seeded into an 8 well PDL-coated plated in Seahorse XF RPMI medium, pH 7.4 (Agilent), supplemented with 10mM glucose and 2mM glutamine. The plate was centrifuged at 200 g for 5 mins and rested in a non-CO2 incubator for 45 mins before analysis. The cartridge and plate were loaded into an Agilent Seahorse XFp Analyzer and cells were injected with succinate (5 mM), oligomycin (1.5 µM), FCCP (0.5 µM) and rotenone + antimycin A (0.5 + 0.5 µM). Data was analysed using Agilent Seahorse Analytics.

#### MDMs

For MDM seahorse analysis, PBMCs were isolated from peripheral blood buffy coats (obtained from the Irish Blood Transfusion Services), from which a monocyte enriched population was obtained by Percoll (Sigma) density gradient centrifugation. 3×10^6^ cells/mL were seeded into an untreated 6 well plate and left to adhere overnight in cRPMI (10% human serum). Cells were washed to further select for monocytes through plastic adherence and differentiated into MDMs for 7 days. MDMs were seeded at a density of 1×10^5^ cells/well in a Seahorse XFe96/XF Pro Cell Culture Microplate at left to adhere overnight. Culture media was replaced by Seahorse XF RPMI medium, pH 7.4 (Agilent), supplemented with 10mM glucose, 2mM glutamine and equilibrated in a non-CO2 incubator for 45 mins before analysis. The cartridge and plate were loaded into an Agilent Seahorse XFe96 Analyzer and cells were injected with oligomycin (1.5 µM), FCCP (0.5 µM) and rotenone + antimycin A (0.5 + 0.5 µM).

## Study approval

Healthy donor bloods were obtained with consent from donors (ethical approval, School of Medicine Research Ethics Committee, Trinity College Dublin). Human bronchoalveolar lavage (BAL) fluid was obtained at bronchoscopy after written informed consent, as approved by the St James’s Hospital (SJH)/ Tallaght University Hospital (TUH) Joint Research Ethics Committee.

## Statistical analysis

GraphPad Prism V.8.00 (GraphPad Software) was used for statistical analysis. Normality was determined using the D’Agostino-Pearson omnibus test. Data were then analyzed using a Student’s t-test when two data sets were being compared, or a one-way/two-way ANOVA test when more than two data sets were being compared. A p value of < 0.05 was considered statistically significant.

## Data availability

All data needed to support the conclusions of the paper are present in the paper or the Supplementary Materials. The BAL single cell RNAseq dataset can be found at^22^.

## Acknowledgments

This research was funded by the Health Research Board of Ireland and by the Royal City of Dublin Hospital Trust. S.M.C is supported by a Research Ireland Future Research Leaders Grant (FRL4862) and a Laureate Award (IRCLA/2022/3619). S.C.D. is supported by Research Ireland. We thank the lab of David Finlay and lab member Carrie Corkish for their advice and guidance on click chemistry.

## Author contributions

Conceptualization, L.G. and K.S.; Methodology, K.S. and L.G.; Software, K.S., A.N., A.S; Formal Analysis, K.S., A.N., A.S., J.M.G.; Investigation, K.S., A.N., A.S., J.M.G., Resources, L.G., J.K., S.M.C., S.C.D.; Writing – Original Draft, K.S. and L.G.; Writing – Review & Editing, K.S., L.G. and S.M.C.; Visualization, K.S., A.N., A.S.; Patient sample collection, O.O.G., L.F., E.M.N., F.O.C., P.N., P.M., S.C.D., J.K., L.G.; Supervision, L.G.

---

**S1.**
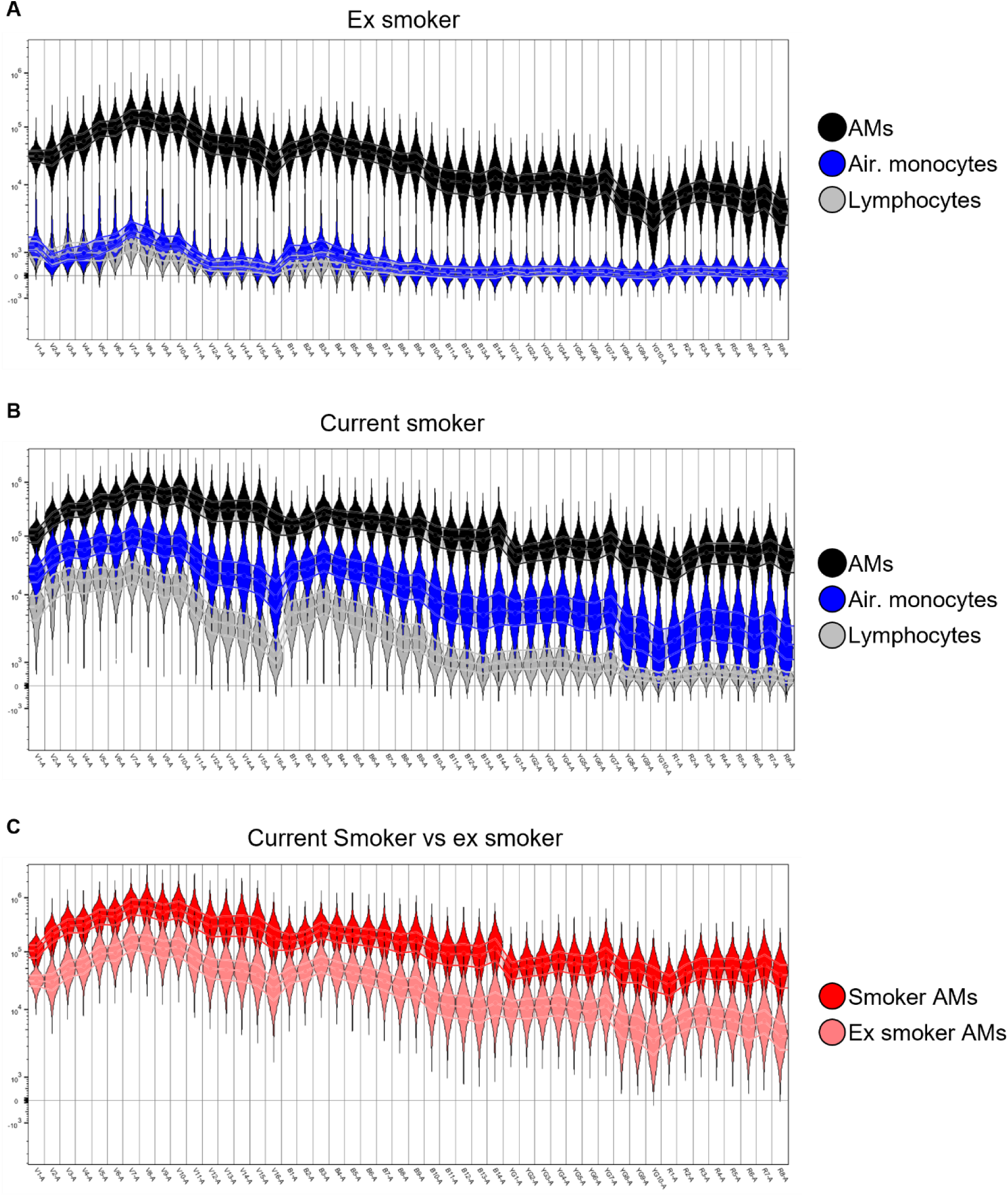
Autofluorescence of *ex vivo* lung leukocytes in ex-smokers and smokers. Cells were isolated from BAL samples, fixed and acquired on a Cytek Aurora flow cytometer. Autofluorescence of *ex vivo* lung leukocytes in (A) ex-smokers and (B) current smokers. (C) Overlay of AM autofluorescence in ex-smokers and current smokers. Air=airspace.

**S2.**
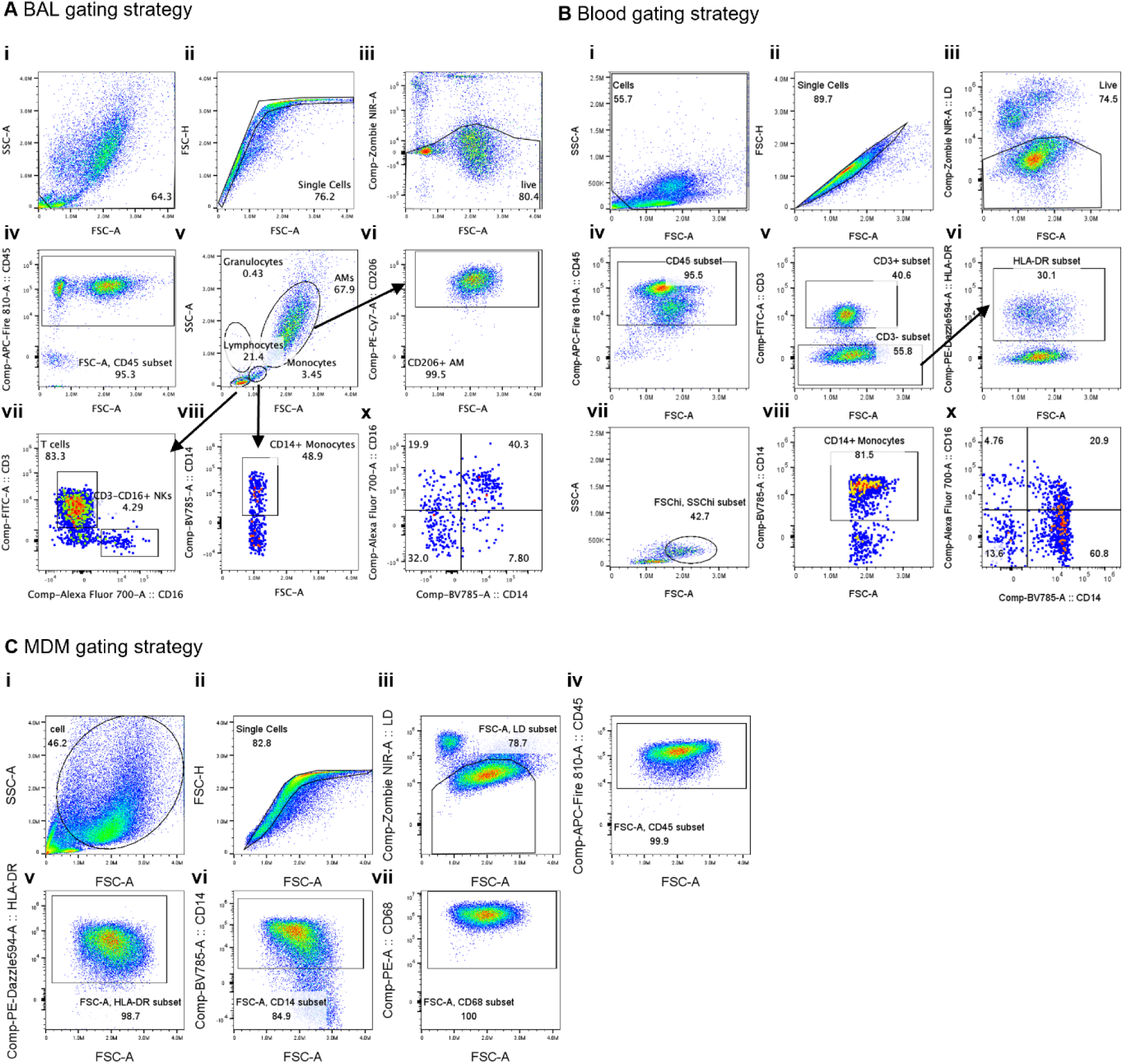
Flow cytometry gating strategies for BAL and blood samples and for MDMs. (A) Gating strategy for BAL samples. Total cells excluding debris were gated on (i) and single (ii), live (iii) and CD45+ cells (iv) were gated on. Lymphocytes, granulocytes, monocytes and AM were first gated on based on FSC and SSC. CD206 identified AM (vi). CD3 identified T cells (vii). CD14 identified monocytes (viii). CD14 and CD16 expression identified monocyte subsets. (B) Gating strategy for blood samples. Total PBMC excluding debris were gated on (i) and single (ii), live (iii) and CD45+ cells (iv) were gated on. Non-T cells were gated on as CD3-(v). HLA-DR+ cells were gated on (vi) and monocytes were identified as FSChi and SSChi to exclude remaining lymphocytes (vii). CD14 identified monocytes (viii). CD14 and CD16 expression identified monocyte subsets (ix). (C) Gating strategy for MDMs. Total cells excluding debris were gated on (i) and single (ii), live (iii) and CD45+ cells (iv) were gated on. MDMs were next characterised as (v) HLA-DR+, (vi) CD14+ and (vii) CD68+.

**S3.**
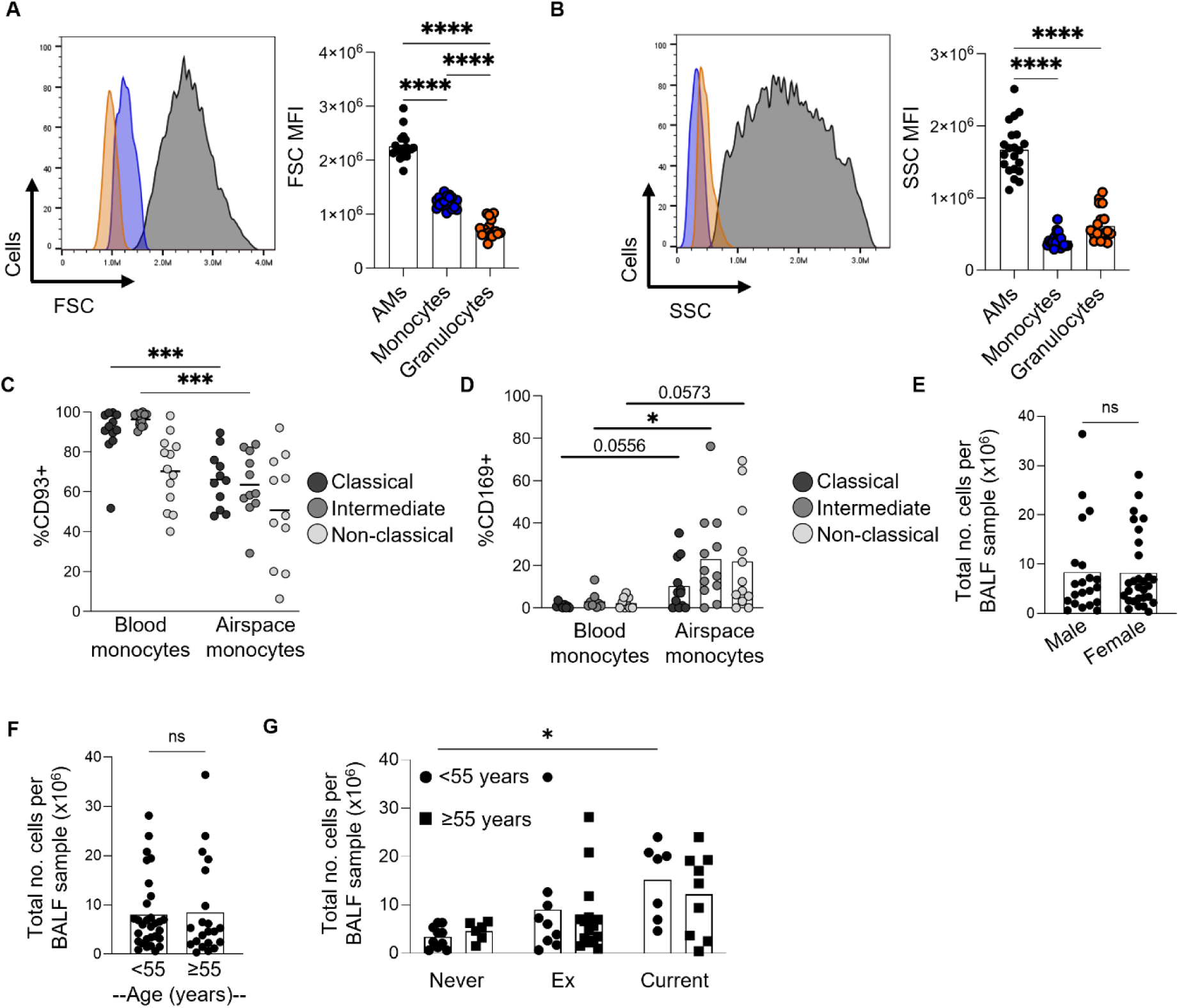
Airspace monocytes have increased expression of CD169 compared to blood monocytes. (A) Representative histogram and pooled data for FSC MFI as a measure of cell size and (B) SSC MFI as a measure of granularity. (C-D) Pooled data of the frequency of CD93 and CD169 expression in monocyte subsets from fresh blood and BAL samples. (E-G) Total cellular yield of BAL samples according to sex and age. (C-D and H) Data was compared using a 2way ANOVA test. (E-F) Data was compared using an unpaired t test. ns = not significant. N = 7-21, *p<0.05, ***p<0.001, ****p<0.0001.

**S4.**
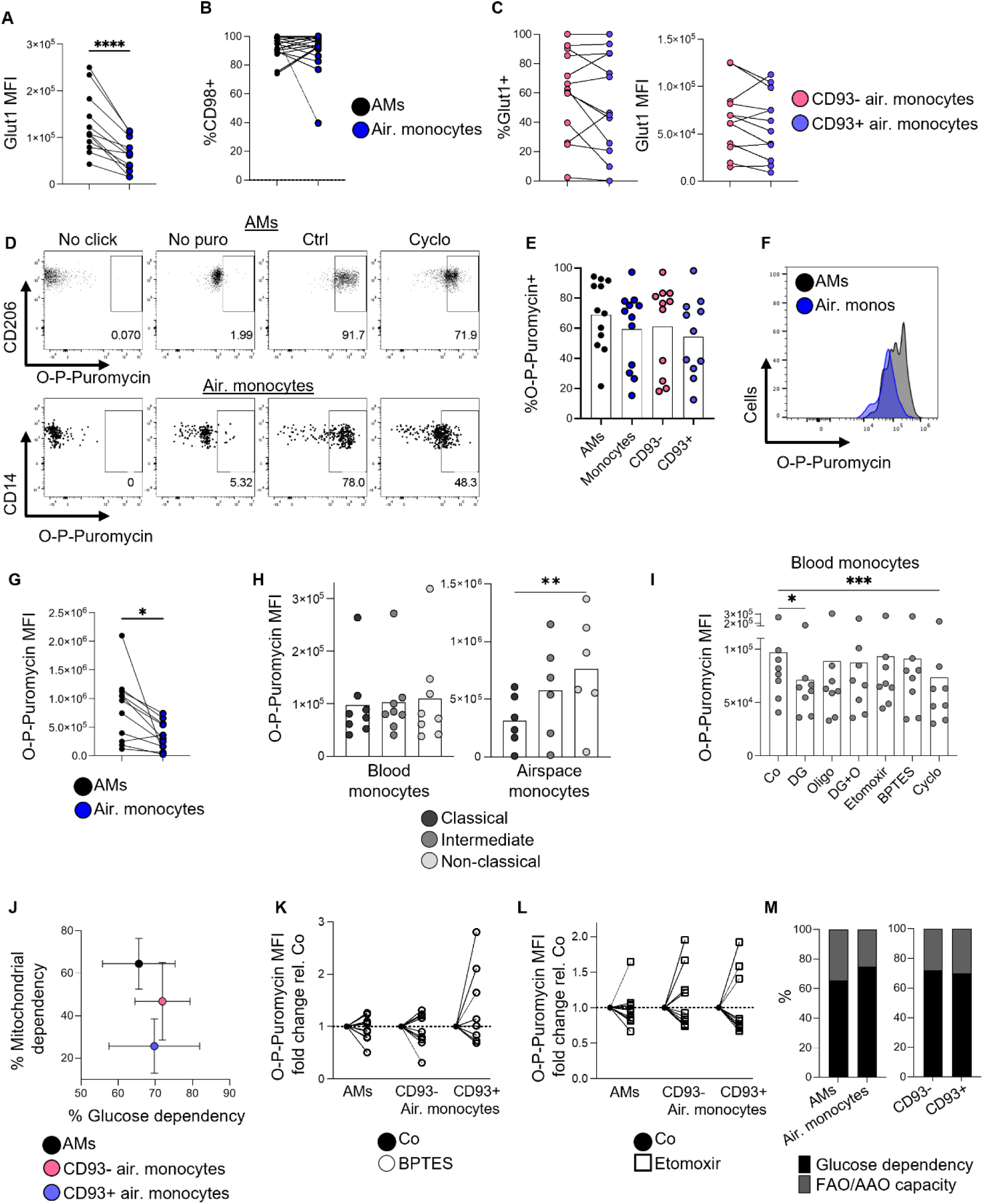
Non-classical monocytes have higher levels of *de novo* protein synthesis than classical monocytes in the human lung. (A-B) Pooled data for Glut1 MFI and frequency of CD98 expression in AM and total airspace monocytes. (C) Pooled data for Glut1 MFI and frequency of Glut1 expression in CD93- and CD93+ airspace monocytes. (D) Representative dot plots for *de novo* protein synthesis assay. Cyclo (cycloheximide) was used as a negative control. (E) Pooled data for the frequency of O-P-Puromycin incorporation into new proteins in AM and airspace monocytes. (F) Representative histogram and pooled data for O-P-Puromycin MFI in AM and airspace monocytes. (G) Pooled data for O-P-Puromycin MFI in AM and airspace monocytes. (H) *De novo* protein synthesis in monocyte subsets from blood and BAL samples. (I) Pooled data of impact of metabolic inhibitors on O-P-Puromycin MFI in blood monocytes. (J) SCENITH metabolic parameters in AM and airspace monocytes. (K-L) Impact of BPTES (5 µM) or etomoxir (5 µM) on protein synthesis in AM and CD93- and CD93+ airspace monocytes. (M) %Glucose dependency and fatty acid/amino acid oxidation capacity (FAO/AAO) in AM, CD93-airspace monocytes and CD93+ airspace monocytes. (A and G) Data was compared using a paired t test. (H-I) Data was compared using a non-parametric One-Way ANOVA test. Air=airspace. N = 6-21, *p<0.05, **p<0.01, ****p<0.0001.

**S5.**
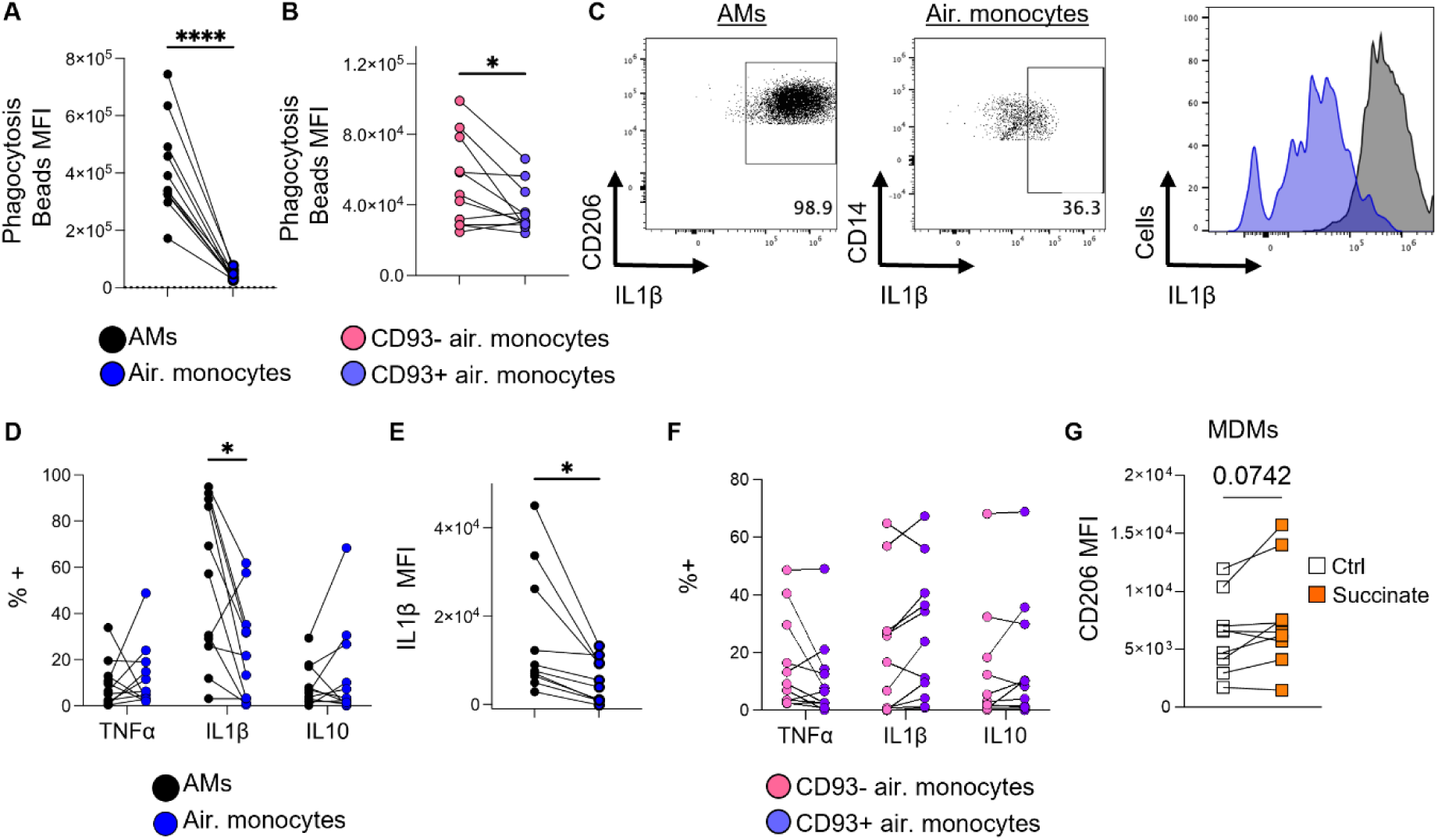
CD93+ airspace monocytes have low phagocytotic capacity. (A-B) *Ex vivo* BAL cells were incubated with FITC-labelled phagocytosis beads for 50 mins and analysed by flow cytometry. Pooled data of phagocytosis bead uptake in AM vs airspace monocytes and in CD93-vs CD93+ airspace monocytes. (C) Representative dot plot and histogram for IL1β expression in AM and airspace monocytes. (D-F) Pooled data of cytokine expression AM and airspace monocytes. (G) CD206 expression in MDMs expanded for 8 days in the presence of absence of succinate (5 mM). (A-B, E and G) Data was compared using a paired t test. Air=airspace. (D and F) Data was compared using a Wilcoxon matched pairs signed rank test. N = 10-11, *p<0.05, ****p<0.0001.

**S6.**
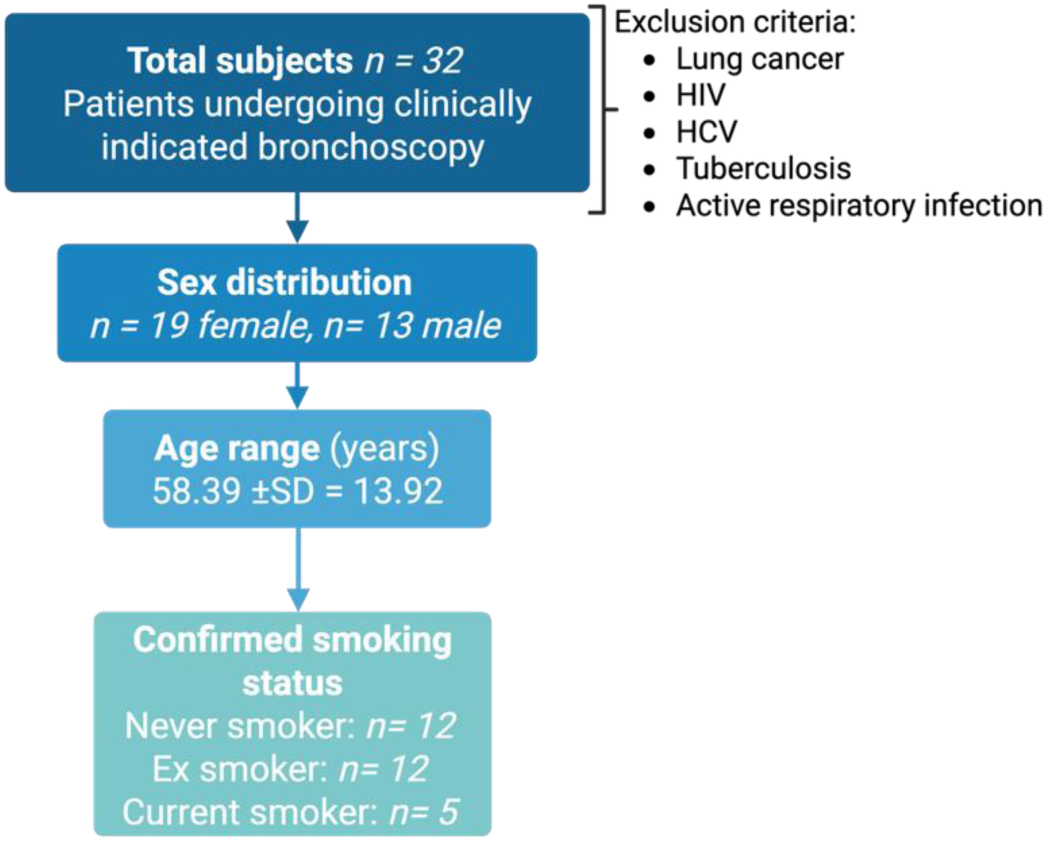
Flow diagram describing the study design and included patients.

